# Disabling Müller Glia Preserves Retinal Function After Retinal Injury

**DOI:** 10.64898/2026.04.14.718211

**Authors:** Daniel Larbi, Shaoheng Chen, Luke D. Gibbons, Ashley Indictor, Seoyoung Kang, Alexander M. Rief, Stefanie G. Wohl

**Affiliations:** Department of Biological and Vision Sciences, The State University of New York, College of Optometry, New York, New York, USA; Indiana University School of Optometry, Bloomington, Indiana, USA; Center for Gene and Cell Therapy, Korea Research Institute of Bioscience and Biotechnology (KIBB), Daejeon, Republic of Korea

**Keywords:** retinal degeneration, Dicer1, light damage, photoreceptors

## Abstract

We developed a physiologically relevant light damage model in pigmented mice and determined how Müller glial (MG) Dicer1 loss impacts retinal structure and function after injury. A moderate light damage paradigm (5,000 lux, 4 hours) was developed in pigmented mice carrying the RPE65 Leu450 variant. MG-specific Dicer1 conditional knockout (cKO) mice across three Cre lines (Rlbp1-CreER, Glast-CreER, Ascl1-CreER) were subjected to light damage at different developmental stages. Retinal structure and function were assessed using optical coherence tomography (OCT), histology, and electroretinography (ERG). Preconditioning and double-damage paradigms were included as controls. The model induced progressive photoreceptor degeneration characterized by early functional decline, followed by structural loss and delayed inner retinal impairment. Across all lines with Dicer loss in MG, retinal structure and function were better preserved following injury than in light-damaged controls. The most sustained protective phenotype was observed in the Rlbp1-CreER-driven line. Inner retinal function (Vmax) was consistently maintained despite reduced photoreceptor input. This phenotype was independent of age, timing of MG manipulation, or baseline retinal condition and was not reproduced by preconditioning paradigms. Dicer-deficient MG displayed reduced glial fibrillary acidic protein (GFAP) immunoreactivity, indicating a potential suppression of glial reactivity. However, the absence of a neuroprotective phenotype following preconditioning suggests that reduced GFAP expression alone is insufficient to account for the observed retinal preservation. Collectively, these findings demonstrate that MG-specific Dicer1 deletion is associated with a neuroprotective retinal phenotype characterized by preserved inner retinal function and reduced secondary degeneration. These findings establish a glia-driven component of retinal degeneration and demonstrate that altering the MG injury response can preserve retinal function following injury.

## Introduction

Retinal degenerative diseases, such as retinitis pigmentosa (RP) and age-related macular degeneration (AMD), are characterized by progressive loss of photoreceptors, the neurons responsible for capturing light and initiating visual processing. Typically, rods are affected first, leading to night blindness and peripheral vision loss, followed by secondary cone degeneration, which impairs central and color vision ^1^. The mechanisms underlying photoreceptor degeneration are complex and involve genetic mutations, oxidative stress, metabolic dysregulation, and inflammation. While late-stage pathology is well described, the early and intermediate processes driving disease progression, particularly the transition from primary photoreceptor damage to secondary neuronal loss, remain incompletely understood.

To study these mechanisms, a range of experimental models have been developed. Genetic models, including rd1, rd10, and P23H mice, have provided important insights into mutation-specific disease pathways (see review ^2^). In parallel, light-induced retinal damage has emerged as a widely used and highly adaptable model to study photoreceptor degeneration and to mimic key features of AMD and RP ^3–8^. While genetic models allow investigation of defined mutations, light damage models offer temporal control and enable the study of environmental, metabolic, and glial contributions to degeneration.

In light damage paradigms, photoreceptor degeneration is typically initiated by excessive rhodopsin activation, leading to metabolic overload, accumulation of reactive oxygen species (ROS), and apoptotic cell death ^8, 9^. Degeneration follows a biphasic pattern, with an early phase of rapid photoreceptor loss and outer nuclear layer thinning, followed by a slower progressive phase ^10^. Rod loss ultimately leads to secondary cone degeneration ^11, 12^ and is accompanied by MG activation, microglial infiltration, and secondary neuronal loss ^1, 2, 5, 8, 11, 13–28^.

Despite their widespread use, many light damage studies rely on albino animals, which are highly susceptible to light exposure ^29–33^ but differ from pigmented retinas in several key aspects. Albino animals lack melanin in the retinal pigment epithelium (RPE), reducing protection against photo-oxidative stress ^34, 35^. They also exhibit altered retinal development, reduced visual function, and elevated baseline MG reactivity ^36–38^, raising concerns about translational relevance.

In contrast, pigmented retinas more closely resemble the human condition, particularly regarding RPE function and oxidative stress regulation. However, commonly used pigmented strains such as C57BL/6 mice are relatively resistant to light damage due to the RPE65 Met450 variant ^39^. This highlights the need for robust and reproducible injury paradigms in pigmented mice that allow controlled induction of degeneration while preserving physiological relevance.

Beyond the need for improved models, there is increasing recognition that retinal degeneration is not solely a photoreceptor-autonomous process. Instead, non-neuronal cells, particularly MG, play a central role in shaping disease outcomes. MG are essential for maintaining retinal homeostasis by providing structural support, regulating ion and neurotransmitter balance, recycling photopigments, and supplying metabolic substrates to neurons (see comprehensive reviews ^18, 40–43^). In response to injury, MG undergo reactive gliosis, characterized by molecular and morphological changes, including upregulation of glial fibrillary acidic protein (GFAP) ^24, 44–50^. While this response can initially support tissue repair, prolonged or excessive glial reactivity has been associated with inflammation, disruption of homeostasis, and secondary neuronal degeneration ^22, 25–27, 41, 42, 46, 51^.

Emerging evidence suggests that regulation of MG reactivity may depend on Dicer1/microRNAs (miRNAs), which control gene expression at the post-transcriptional level and regulate entire gene networks. They have been implicated in retinal development ^52–59^, homeostasis ^60–64^, and responses to injury or disease ^65–75^. Notably, depletion of miRNAs in MG via Dicer1 deletion leads to retinal degeneration without GFAP upregulation, suggesting that miRNAs are required for glial reactivity ^60^.

However, how retinas with Dicer1-depleted MG respond to acute injury has not been explored. In the present study, we established a moderate light damage model that enables controlled induction of photoreceptor damage in pigmented mice with MG-specific Dicer1 deletion. A particular emphasis was placed on *in vivo* assessments of retinal structure and function *in vivo*, using clinically relevant techniques: spectral domain optical coherence tomography (OCT) and electroretinography (ERG). Since retinal function is the most clinically relevant measure and does not always correlate with structural or molecular changes, we compared multiple models and injury paradigms.

We hypothesized that Dicer1 is a critical regulator of the Müller glial injury response and that its depletion alters disease progression, resulting in measurable changes in retinal structure and function assessed by OCT and ERG. By combining a physiologically relevant light damage model with inducible, MG-specific Dicer1 deletion, this study establishes a glia-driven component of retinal degeneration and provides a foundation for future studies aimed at identifying the Dicer1-dependent molecular mechanisms that underlie this neuroprotective phenotype.

## Methods

### Transgenic mice and Cre induction

All experiments were conducted at the State University of New York, College of Optometry, in accordance with Institutional Animal Care and Use Committee (IACUC) protocols and the ARVO Statement for the Use of Animals in Ophthalmic and Vision Research.

To label and manipulate MG, *Rlbp1-CreERT2* ^76^ and *Glast-CreERT* (Tg[Slc1a3-cre/ERT]1Nat, ID 012586, Jackson Laboratories) strains were used, while the *Ascl1-CreERT2* line (ID 12882 ^77^, Jackson Laboratories) was employed to target MG progenitors/precursors and young MG. These Cre lines were crossed with the *R26-stop-flox-CAG-tdTomato* strain (Ai14, #007908, Jackson Laboratories) to generate the wildtype (wt) reporter mice, and with the Dicer1 conditional knockout strain (*Dicerf/f*, #006001 ^78^, Jackson Laboratories) to generate the conditional Dicer1 knockout strains (Dicer-cKO). All lines were additionally crossed with the S129 strain (129S1/SvImJ, #002448, Jackson Laboratories), which carries the RPE65 Leu450 variant conferring susceptibility to light damage ^79, 80^. Genotyping primers are listed in Table S1. Cre specificity had been previously validated ^57, 61, 65^.

Strains are referred to as follows:: *Rlbp1-CreER:stop^f/f^-tdTomato:RPE65^450Leu^*(wildtype or Rwt), *Glast-CreER:stop^f/f^-tdTomato:RPE65^450Leu^*(wildtype or Gwt), *Ascl1-CreER:stop^f/f^-tdTomato:RPE65^450Leu^*(wildtype or Rwt); *Rlbp1-CreER:stop^f/f^-tdTomato:RPE65^450Leu^*:Dicer*^f/f^*(Rlbp-Dicer cKO or RcKO), *Glast-CreER:stop^f/f^-tdTomato:RPE65^450Leu^*:Dicer*^f/f^*(Glast-Dicer cKO or GcKO), *Ascl1-CreER:stop^f/f^-tdTomato:RPE65^450Leu^*:Dicer*^f/f^*(Ascl1-Dicer cKO or AcKO). We also generated Cre-negative *stop^f/f^-tdTomato:RPE65^450Leu^ and stop^f/f^-tdTomato:RPE65^450Leu^*:Dicer*^f/f^*.

CreERT recombination was induced by intraperitoneal tamoxifen (75 mg/kg, Sigma-Aldrich, St. Louis, MO) in corn oil for four consecutive days at postnatal days P0–3 (Ascl1-Cre) or P11–14 (Rlbp1-Cre, Glast-Cre). Control groups included S129 and Cre-negative mice and were compared to tamoxifen-treated wild-type animals to control for tamoxifen or background strain effects. Both sexes were used. Mice with insufficient Cre-induction were excluded.

### Light damage

Mice were dark-adapted overnight to enhance retinal sensitivity. Prior to light exposure, pupils were dilated with 1.0% tropicamide. Animals were then exposed to diffuse, cool white light at 5,000 lux for 4 hours in a home-made setup ^65^ following instructions from Grimm and Reme ^3^. Food and gel remained available throughout exposure. Light intensity was measured at cage level using a lux meter, and ambient temperature was maintained at ∼25–27°C. Following exposure, mice were returned to standard housing under a 12-hour light/dark cycle. Retinal analyses were performed at 1, 2, 3, 7, 14, and 28 days post-exposure.

For preconditioning, mice were exposed to 1,000 lux for 2 hours, followed by a second exposure at day 3 to 5,000 lux for 4 hours. For the double-damage paradigm, mice received 5,000 lux for 4 hours, followed by a second identical exposure on day 3. In both paradigms, animals were analyzed 7 days after the 5000-lux exposure.

### Retinal spectral-domain optical coherence tomography (OCT) imaging

To assess retinal structure in vivo, OCT was performed using the Envisu R2200 system (Bioptigen, Durham, NC), as described previously ^57, 61^. Mice were anesthetized with ketamine (75–100 mg/kg) and xylazine (5–10 mg/kg) in sterile saline, and pupils were dilated with 2.5% phenylephrine hydrochloride and 1.0% tropicamide. Corneal hydration was maintained with 0.5% carboxymethylcellulose sodium during imaging. Rectangular and radial volume scans (1.4 × 1.4 mm; 1000 A-scans per B-scan × 15 frames per B-scan) centered on the optic nerve head were acquired.

Retinal layers were manually segmented using DiverRelease_2_4 software within a 9 × 9 grid across nasal–temporal and superior–inferior axes. Total retinal thickness was defined from the inner border of the nerve fiber layer (NFL) to the outer border of the retinal pigment epithelium (RPE). While all layers were quantified, the analysis focused on the outer retina. Due to limited resolution after damage, the region between the outer limiting membrane (OLM) and RPE was analyzed as a combined OLM–RPE layer ^81, 82^. Measurements were obtained from the central retina (∼650 μm radius) and averaged across eccentricities and quadrants for each time point. Groups included mixed sexes with ≥4 mice per group. Data are presented as mean ± standard deviation (SD).

### Electroretinogram (ERG) recordings

Full-field ERGs were performed as previously described ^57, 61^. Mice were dark-adapted overnight and anesthetized with ketamine (75–100 mg/kg) and xylazine (5–10 mg/kg) in sterile saline. Recordings were conducted in a dark room using the Espion system (Diagnosys LLC, Lowell, MA, USA). Pupils were dilated with 1% tropicamide, and corneal hydration was maintained with 0.5% carboxymethylcellulose sodium. Body temperature was stabilized using a heating pad. A gold wire electrode was placed on the cornea, with subdermal reference and ground electrodes positioned in the cheek and back, respectively.

Full-field flashes (5 ms) were delivered using a handheld stimulator, and scotopic ERGs were recorded across 17 light intensities (0.001–64 cd·s/m²). A-wave amplitudes were measured from baseline to the trough of the initial negative deflection, and b-wave amplitudes from this trough to the peak of the subsequent positive deflection. Scotopic b-wave intensity–response data were fitted using the Naka–Rushton equation:

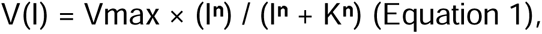

where Vmax represents maximal response amplitude, K the semi-saturation constant, and n the slope ^83, 84^.

Groups included mixed sexes with ≥4 mice per group (preconditioning group n = 3). Data are presented as mean ± standard error of the mean (SEM).

### Tissue preparation and Immunofluorescent labeling

Mice were euthanized, and their eyes were marked nasally for orientation, enucleated, and transferred to cold phosphate-buffered saline (PBS). Eyeballs were fixed in 4% paraformaldehyde (PFA) at 4°C for 20 minutes. After removal of the cornea, lens, iris, and vitreous, eyecups were post-fixed in 4% PFA for 20 minutes, washed in PBS, and cryoprotected overnight in 30% sucrose at 4°C. Tissue was embedded in embedding medium - defined orientation, frozen at −80°C, and sectioned at 12 μm.

For immunofluorescent labeling, sections were dried (37°C, 20 min), fixed in 4% PFA, washed, and blocked in 5% horse serum with 0.5% Triton X-100 for ≥1 hour at room temperature. Sections were incubated overnight with primary antibodies (mouse anti-M-Opsin, Sigma/Millipore, AB5405, 1:500; rabbit anti-GFAP, Dako, Z033401-2, 1:1000) dissolved in 5% horse serum solution. After three PBS washes, incubation with secondary antibodies for 1 hour followed (AffiniPure F(ab’)₂ fragment donkey anti-mouse or rabbit IgG (H+L) with Alexa Fluor 488 [715-546-150] or Alexa Fluor 647 conjugation [711-606-152], 1:500, Jackson ImmunoResearch Laboratories, Inc.). Nuclei were counterstained with DAPI (1:1000, Sigma/Millipore), and sections were mounted using Invitrogen mounting medium.

### Confocal laser scanning microscopy, image processing, and analysis

Immunolabeled sections were imaged using a confocal microscope (Olympus FM1200 Fluoview, 40× objective). Images were processed using Adobe Bridge, Photoshop (Adobe, San Jose, CA), and Affinity (Canva, Sydney, AU). For photoreceptor row counts, damaged regions were divided into eight equal segments, and DAPI-positive rows were quantified and averaged across up to four images per animal. Groups included mixed sexes with ≥4 mice. Data are presented as mean ± SD.

### Data analysis and statistics

Analyses were performed in a blinded manner. Statistical analyses were performed using GraphPad Prism 11 and R, with significance defined as *p* < 0.05. Depending on the experimental design, i.e., multiple time points and/or genotypes, datasets were analyzed using a two-way mixed-effects model, one-way ANOVA, or two-way ANOVA, followed by Dunnett’s, Tukey’s, or Bonferroni’s post-hoc multiple comparisons corrections. The specific statistical test, post-hoc correction, and sample sizes for each experiment are explicitly indicated in the corresponding figure legends.

### Artificial Intelligence (AI) tools

Generative AI (ChatGPT, OpenAI) was used to assist with the generation of the summary Figure 12 schematic based on uploaded PowerPoint drafts. Grammarly and ChatGPT were used for language editing of the manuscript text. All AI content was critically reviewed and approved by the authors.

## Results

### Establishment and characterization of a light damage model for moderate photoreceptor degeneration

To induce light damage in pigmented transgenic strains (predominantly C57BL/6 background), mice were crossed with the S129 strain carrying the leucine variant of RPE65, which enhances susceptibility to light damage ^79, 80^. To allow moderate and progressive photoreceptor degeneration suitable for studying MG contributions to neuronal function, a paradigm of 5,000 lux for 4 hours was selected. Light damage was performed in 2-4-month-old mice (mixed strains).

To characterize this model, *in vivo* retinal imaging using OCT was conducted at 7, 14, and 28 days after light damage (dLD; Figure 1A). Undamaged retinas displayed intact layering without abnormalities (Figure 1B), consistent with previous reports ^85, 86^. Total retinal thickness in naïve mice (including all Cre strains and naïve S129 mice) was ∼230 μm (Figures 1C-D), in agreement with earlier studies ^87, 88^. Following light damage, retinal thickness decreased by ∼20% at 7 dLD, 26% at 14 dLD, and 30% at 28 dLD (Figures 1B-D).

**Figure 1:**
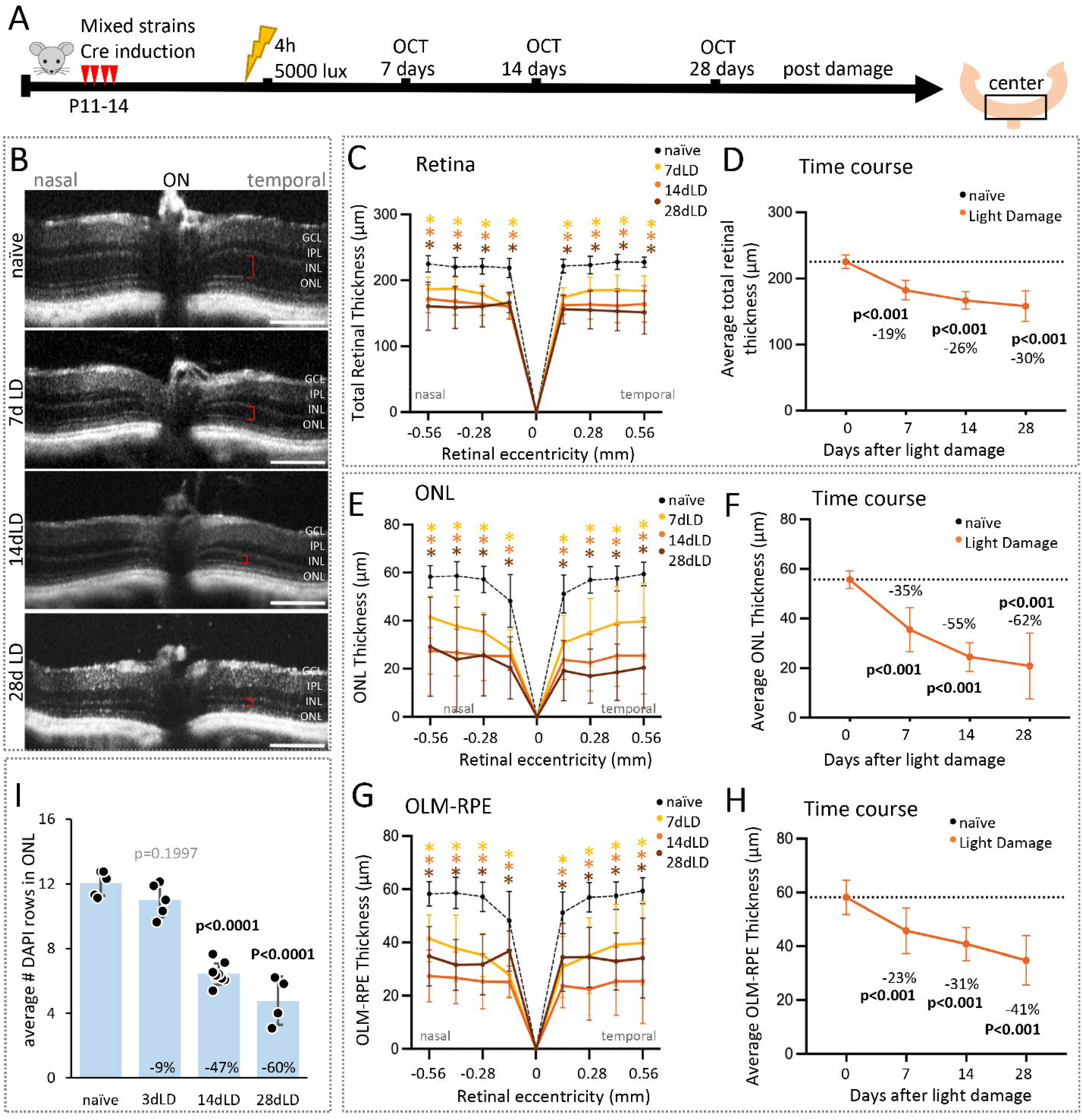
Light damage at 5000 lux for 4h leads to moderate neuronal loss. A: Experimental design of the Optical Coherence Tomography (OCT) series. B: OCT images of center retinas (1300 μm diameter) at the nasal-temporal axis of undamaged or light-damaged mice 7, 14, or 28 days after light damage (dLD). Red brackets indicate ONL thickness. C-H: Spider plots (nasal-temporal axis) and time courses (averaged nasal-temporal and superior-inferior axis) of the thickness (diameter, μm) of the total retina (C, D), the outer nuclear layer (ONL, E, F), and the outer limiting membrane/retinal pigment epithelium (OLM-RPE, G, H) of undamaged (n=11) or light-damaged mice, 7d (n=8), 14d (n=12), or 28dLD (n=6). I: Photoreceptor rows (DAPI counts) in the outer nuclear layer of undamaged (n=5) or light-damaged mice, 3d (n=5), 14d (n=9), or 28dLD (n=6). Mean ± S.D., significant differences between LD and undamaged control are indicated using two-way mixed-effects model with Dunnett’s multiple comparisons correction comparing all time points to the naïve control (C, E, G: *: p≤0.05) and linear mixed-effects model with post hoc comparisons using estimated marginal means and Bonferroni correction for multiple comparisons; p-values for time-line graphs are given in the graphs. Scale bars in B: 200 µm. GCL: ganglion cell layer, IPL: inner plexiform layer, INL: inner nuclear layer, OPL: outer plexiform layer, ONL: outer nuclear layer, OLM: outer limiting membrane.

Given that photoreceptors are the primary target of light damage, we focused on the ONL and the photoreceptor segment region between OLM and RPE. In undamaged mice, ONL thickness was ∼56 μm. After light damage, the ONL declined successively by ∼35%, ∼55%, and ∼62%, 7d, 14d, and 28d, respectively; (naïve: 56 μm, 7dLD: 36 μm, p<0.001; 14dpLD: 25 μm, p<0.001; 28dpLD: 21 μm, p<0.001, Figures 1B, E-F).

Similarly, the OLM-RPE region (∼59–60 μm in controls) decreased by ∼23%, ∼31% and ∼41%, 7d, 14d, and 28d, respectively; (naïve: 59 μm, 7dpLD: 46 μm, p<0.001; 14dpLD: 41 μm, p<0.001; 28dpLD: 35 μm, p<0.001, Figures 1B, G-H). To validate these *in vivo* findings, photoreceptor rows (DAPI+) were quantified in retinal sections. We included a 3-day time point at which no significant loss was detected. At 14 and 28dpLD, cell numbers were reduced by ∼47% and ∼60%, respectively (Figure 1I), consistent with our OCT measurements (55% and 62%, respectively).

These data indicate biphasic degeneration kinetics, with approximately half of the photoreceptors lost within the first two weeks, followed by a plateau phase. Notably, compared to previous models, this paradigm results in more moderate degeneration, with ∼40% of rods remaining at 28 dpLD. Inner retinal layers were largely preserved at these time points (Figures S1A,B).

We next assessed retinal function using full-field ERG (Figure 2A). Scotopic ERGs were performed to evaluate rod, rod-bipolar, and MG function. In undamaged mice, a-wave amplitudes (rod function) were ∼230 μV (Figures S1C-D). Following light damage, a-wave amplitudes decreased by ∼46%, ∼55%, and ∼56% at 7d, 14d, and 28d pLD respectively (naïve: 227 μV, 7dpLD: 121 μV, p<0.001; 14dpLD: 100 μV, p<0.001; 28dpLD: 99 μV, p<0.001, Figure 2B). Time course analysis confirmed a biphasic decline similar to structural changes (Figure 2C). Notably, at 3d pLD, a-wave amplitudes were already reduced by ∼30% despite no detectable structural loss (Figure S1E). This indicated that functional decline precedes photoreceptor degeneration, consistent with disease progression ^11^.

**Figure 2:**
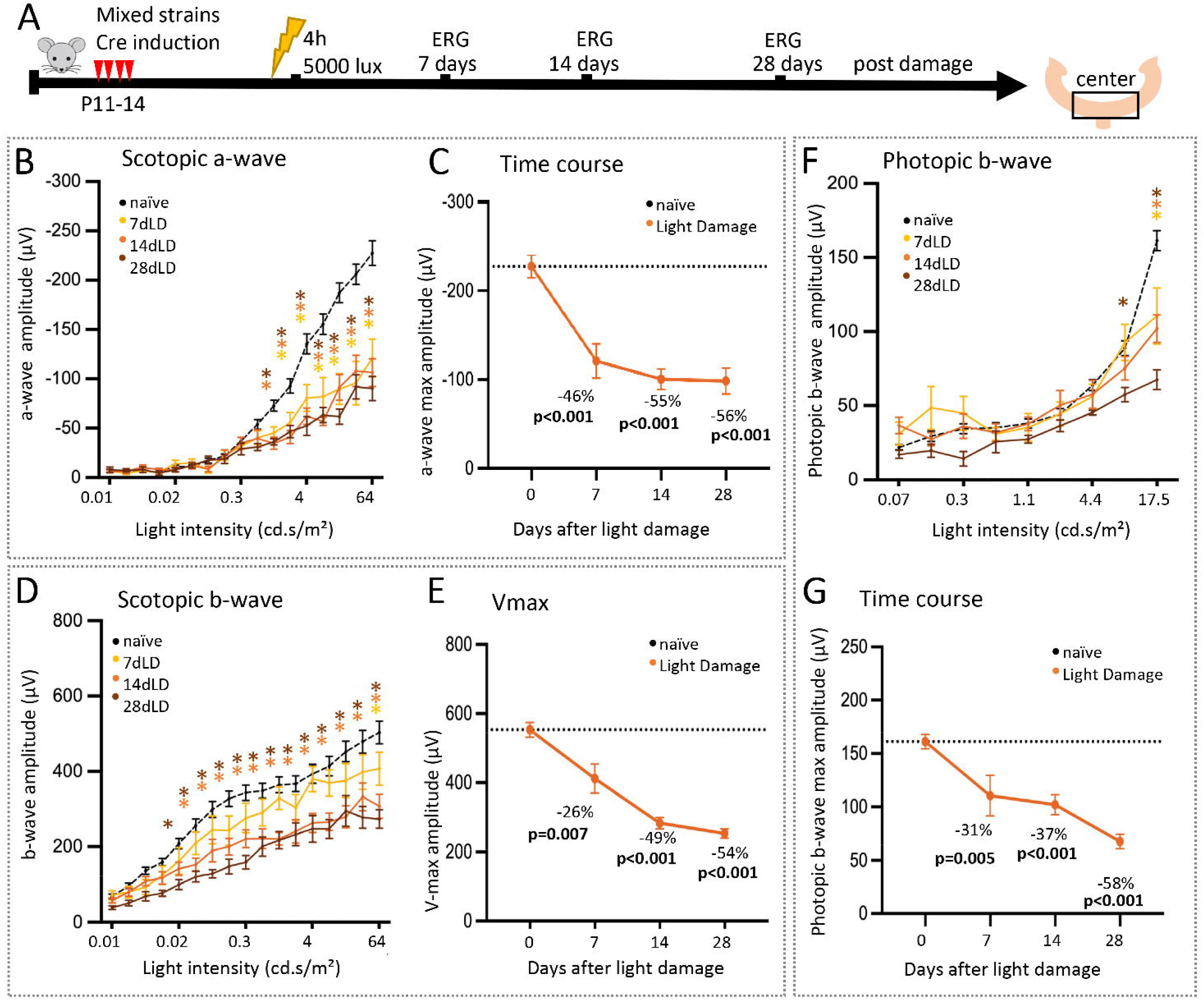
Light damage at 5000 lux for 4h leads to moderate loss of inner retinal function. A: Experimental design of the electroretinography (ERG) series. B-G: Full-field ERG recordings showing scotopic a-wave amplitudes (B) and corresponding time line of the averaged amplitudes (C), scotopic b-wave amplitudes (D) and estimated saturated amplitudes (Vmax, responsiveness) using the Naka-Rushton equation (E), as well as photopic b-wave amplitudes (F) and corresponding time line of the averaged amplitudes (G) of undamaged (n=27) or light-damaged mice, 7d (n=7), 14d (n=9), or 28dLD (n=6). Mean ± S.E.M. Significant differences between LD and undamaged mice are indicated using a two-way mixed-effects model with Dunnett’s multiple comparisons correction comparing all time points to the naïve control (C, E, G: *: p≤0.05) and a linear mixed-effects model with post hoc comparisons using estimated marginal means and Bonferroni correction for multiple comparisons; p-values for time-line graphs are given in the graphs.

Rod bipolar cell and MG function were evaluated by the scotopic b-wave amplitudes, which usually follow the input patterns (lower a-wave results in lower b-wave). Scotopic b-wave amplitudes (∼500 μV in controls) showed a delayed decline, decreasing by ∼20% at 7 dLD, ∼40% at 14 dLD, and ∼46% at 28 dpLD (naïve: 503 μV, 7dpLD: 408 μV, p=0.1445; 14dpLD: 309 μV, p<0.0001; 28dpLD: 273 μV, p=0.0002. Hence, the b-wave followed the reduced rod input (a-wave, Figure 2D).

To further characterize inner retinal function, saturated amplitudes (Vmax) were calculated using the Naka–Rushton equation, an established method we used in previous studies to evaluate inner retinal function ^57, 61^. It is a measure of the retina’s total power output or maximum working capacity. A high or normal Vmax means the inner retina cells (rod bipolar cells and MG) are healthy, numerous, and capable of generating a strong response to light. Conversely, a reduced function suggested damaged or a reduced number of cells, lowering the maximum output the retina can produce ^84, 89^. After damage, over time we found a Vmax reduction of ∼26%, ∼50% and ∼54%, 7d, 14d, and 28d pLD, respectively (naïve: 553 μV; 7dpLD: 412 μV, p=0.007; 14dpLD:284 μV, p<0.001; 28dpLD: 253 μV, p<0.001, Figure 2E). Notably, this early reduction was not apparent in raw b-wave amplitudes, indicating that Vmax provides a more sensitive measure of early inner retinal dysfunction.

Since rod loss affects cone health ^10, 11, 13^, we next evaluated the photopic b-wave amplitudes. In undamaged animals, the average amplitude was approximately 160 μV. Following light damage, amplitudes decreased by ∼30%, ∼40 and ∼60%, 7d, 14d, and 28d pLD, respectively (naïve: 160 μV; 7dpLD: 111 μV, p=0.005; 14dpLD: 102 μV, p<0.001; 28dpLD: 68 μV, p<0.001, Figure 2F, timeline in G). This indicates progressive cone dysfunction in parallel with rod degeneration (∼60%, respectively).

In summary, scotopic ERG recordings revealed an early and significant decline in rod function, followed by a delayed reduction in inner retinal responses. Notably, Vmax was already reduced at 7 dLD, indicating early impairment of inner retinal function. Together, these findings establish a moderate light damage model for pigmented transgenic mice that produces sustained, progressive structural and functional degeneration while preserving sufficient retinal integrity for mechanistic studies.

### Retinas with Dicer1-depleted MG display reduced neurodegeneration after light damage

We previously demonstrated that Dicer1-dependent pathways are essential for maintaining normal Müller glial (MG) function and retinal homeostasis ^60, 61^. Furthermore, we showed that miRNA expression is dynamically regulated in FACS-isolated reactive MG following light-induced retinal injury using the same MG reporter mouse ^65^. In addition, Dicer1-depleted MG failed to upregulate glial fibrillary acidic protein (GFAP)^60, 65^, a canonical marker of reactive gliosis, even during advanced retinal degeneration. Because sustained GFAP expression has been associated with maladaptive glial reactivity and neurotoxicity ^24, 44, 46–48, 90, 91^, we hypothesized that Dicer1-depleted MG would exhibit an altered injury response that influences retinal degeneration.

To test this hypothesis, we quantified retinal structure and function longitudinally using OCT and ERG. Adult MG-specific Dicer1 conditional knockout (cKO) mice, previously established and characterized by our group ^60, 61, 65^, were subjected to our moderate light damage paradigm using two independent MG-specific Cre driver lines,the Rlbp1-CreER: Dicer^f/f^: tdTomato^stopf/f^ and the Glast-CreER: Dicerf/f: tdTomato^stopf/f 61^. Both lines mediate Cre recombination in approximately 80–95% of MG following tamoxifen induction ^60, 61, 65, 92^.

For the present study, both MG-specific Dicer1-cKO lines were crossed onto the RPE65^450Leu^ background to establish a reproducible light damage model in pigmented mice. Structural, functional, and histological outcomes were subsequently evaluated in light-damaged Dicer1-cKO mice and their corresponding light-damaged reporter controls (Rlbp1-CreER: tdTomato^stopf/f^: RPE65^450Leu^ and Glast-CreER: tdTomato^stopf/f^: RPE65^450Leu^). In both models, Dicer1 deletion was induced by tamoxifen administration at postnatal days 11–14, followed by light damage at 3 months of age.

Successful injury induction was first verified 7 days after light damage (7d pLD), a time point selected to confirm consistent retinal damage and exclude animals with insufficient light exposure. Because light damage may be spatially heterogeneous and not fully detected by OCT, ERG was additionally performed to verify functional impairment. Following confirmation of successful injury induction, the primary analyses of retinal structure and function were conducted at 14 dLD, with an additional assessment at 28 dLD to determine whether the observed phenotype was sustained over time.

We first characterized the response in the Rlbp1-CreER model (Figure 3A), our primary and most extensively validated MG-specific Dicer1-cKO line, which has served as the basis for our previous studies ^60, 61, 65, 92^. Key findings were subsequently examined in the independent Glast-CreER model to assess their reproducibility across distinct MG-specific Cre driver lines.

**Figure 3:**
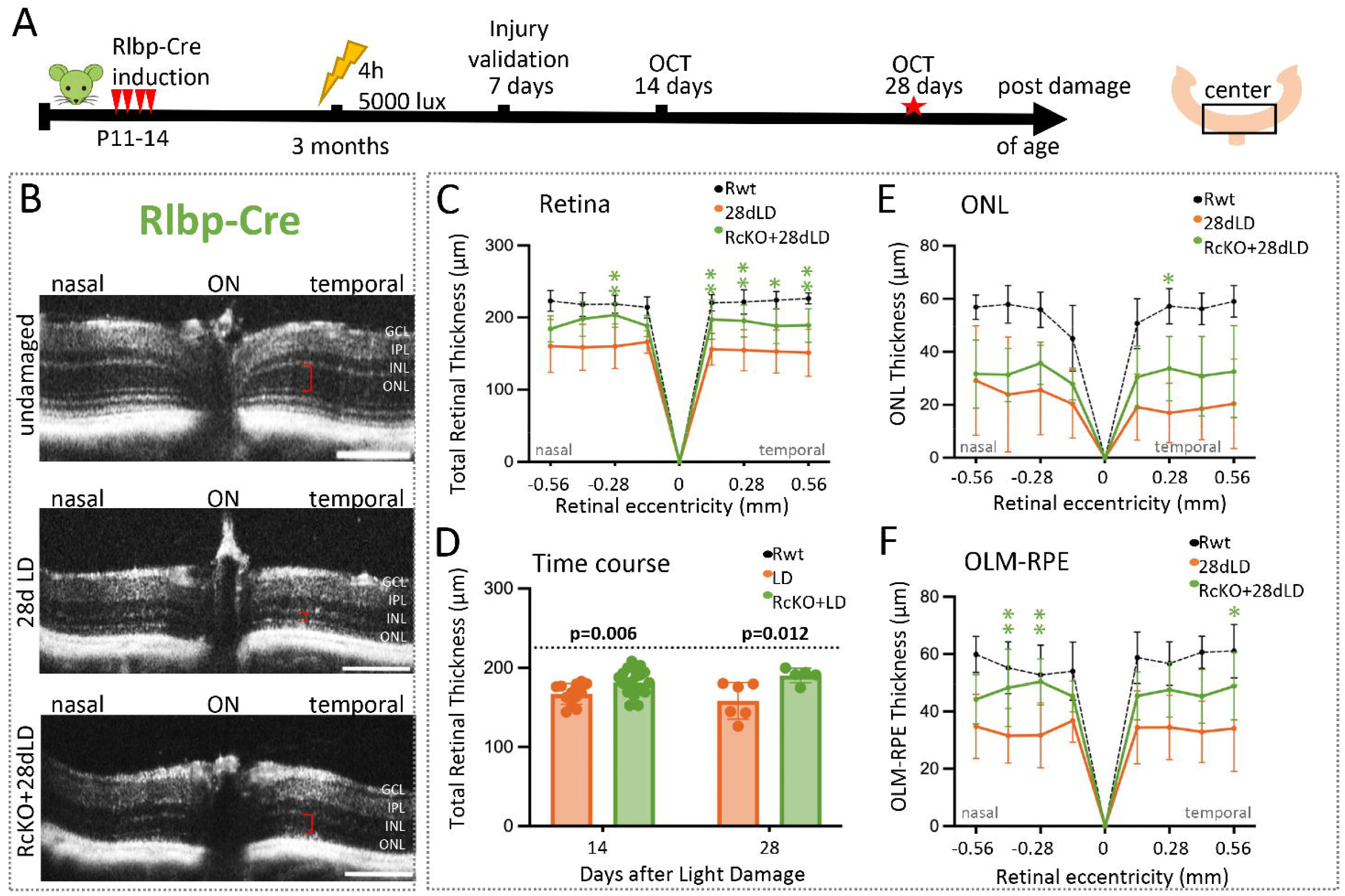
Light-damaged Rlbp-Cre: Dicer-cKO mice display a structurally preserved retina. **A:** Experimental design of the Optical Coherence Tomography (OCT) series. **B:** OCT images of center retinas (1300 μm diameter) at the nasal-temporal axis of undamaged Rlbp-Cre wildtype mice, light-damaged Rlbp-Cre: Dicer-cKO (RcKO+LD), as well as light-damaged wildtypes (LD) 28 days after light damage (dLD). Red brackets indicate ONL thickness. **C-F:** Spider plots (C,E,F, nasal-temporal axis) and time course (D, averaged nasal-temporal and superior-inferior axis) of the thickness (diameter, μm) of the total retina (retina, C, D), the outer nuclear layer (ONL, E), and the outer limiting membrane/retinal pigment epithelium (OLM-RPE, F) of undamaged (n=11), light-damaged Rlbp-Cre: Dicer-cKO (RcKO+LD, n=6) as well as light-damaged wildtypes (LD, n=6), 28 days after light damage (dLD). Mean ± S.D., Significant differences between damaged wildtypes and cKOs are indicated, using a two-way ANOVA with Tukey’s multiple comparisons correction; *: p≤0.05, **: p≤0.01 (C, E, G) and linear mixed-effects model with post hoc comparisons using estimated marginal means and Bonferroni correction for multiple comparisons in D; p-values for time-line graphs are given in the graphs. Undamaged wildtype values serve only as a reference /baseline (dotted lines). Scale bars in B: 200 µm. GCL: ganglion cell layer, IPL: inner plexiform layer, INL: inner nuclear layer, OPL: outer plexiform layer, ONL: outer nuclear layer, OLM: outer limiting membrane.

OCT imaging confirmed photoreceptor loss in Rlbp-Cre: Dicer-cKO retinas as early as 7d pLD, confirming effective damage induction (Figure S2A-C). However, while light-damaged wildtypes exhibited progressive thinning over time (Figures 1C-D, 3B), Rlbp-cre-Dicer-cKO retinas showed relative structural preservation, particularly at 28d pLD. Averaged measurements revealed ∼20% preservation of total retinal thickness at 28 dpLD (LD: 158 μm vs. cKO+LD: 190 μm, p=0.012; Figures 3C-D). Analysis of the outer retina (ONL and OLM–RPE) showed some tendency toward preservation in Rlbp-cre: Dicer-cKO mice (Figures 3E, F, S2D-E), while inner retinal layers remained largely unchanged (Figures S2F-G).

We next assessed retinal function using ERG, first to confirm successful damage at 7d and then to evaluate retinal function during retinal degeneration (Figure 4A). Seven days after LD, ERG recordings further confirmed successful injury; however, it was less severe than in wildtypes (Figures S3A-C). The analysis of scotopic a-wave amplitudes (rod function) 14d and 28d post LD showed significantly higher values in cKO mice compared to light-damaged wildtypes (14 dpLD: 168 μV vs. 100 μV, p=0.012; 28 dpLD: 154 μV vs. 99 μV, p=0.011; Figures 4B-C, S3D).

**Figure 4:**
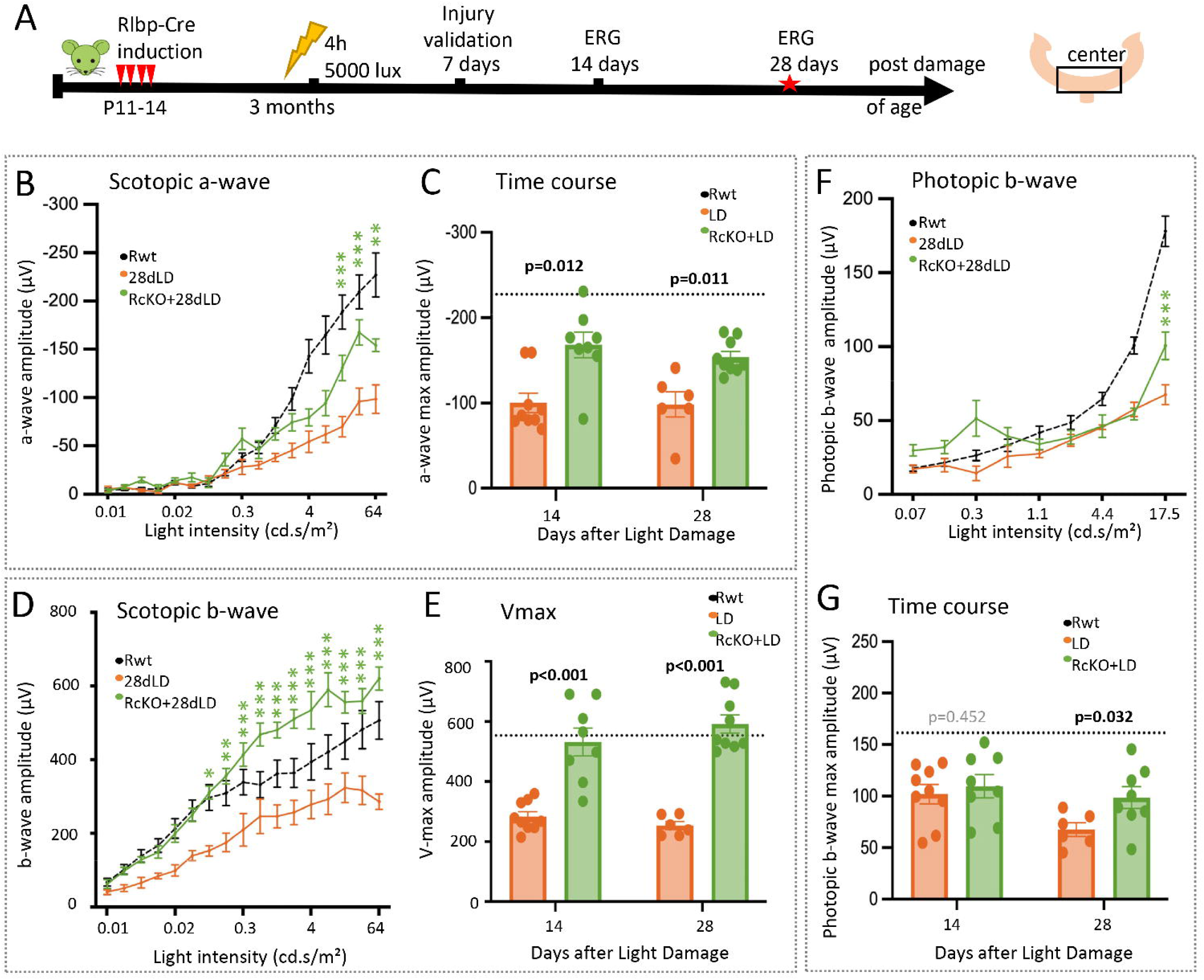
Light-damaged Rlbp-Cre: Dicer-cKO mice have preserved retinal function. A: Experimental design of the electroretinography (ERG) series. B-G: Full-field ERG recordings showing scotopic a-wave amplitudes (B) and corresponding time line of the averaged amplitudes (C), scotopic b-wave amplitudes (D) and estimated saturated amplitudes (Vmax, responsiveness) using the Naka-Rushton equation (E), as well as photopic b-wave amplitudes (F) and corresponding time line of the averaged amplitudes (G) of undamaged Rlbp-Cre wildtypes (Rwt), n=12), light-damaged Rlbp-Cre: Dicer-cKOs (RcKO+LD) 7d (n=7), 14d (n=8), or 28dLD (n=5) as well as light damaged wildtypes (LD) 7d (n=7), 14d (n=7), or 28dLD (n=5). Mean ± S.E.M. Significant differences between damaged wildtypes and cKOs are indicated, using a two-way ANOVA with Tukey’s multiple comparisons correction; *: p≤0.05, **: p≤0.01 (B, D, F) and linear mixed-effects model with post hoc comparisons using estimated marginal means and Bonferroni correction for multiple comparisons in C, E, G; p-values for time-line graphs are given in the graphs. Undamaged wildtype values serve only as a reference /baseline (dotted lines).

Strikingly, scotopic b-wave amplitudes (rod bipolar and MG function) were fully preserved in cKO mice across all time points (14 dLD: 536 μV; 28 dLD: 620 μV), comparable to undamaged controls (∼510 μV; Figures 4D-E, S3E). Consistent with this, Vmax values derived from Naka-Rushton fits were significantly higher in cKO mice compared to light-damaged wildtypes (14 dLD: p<0.001; 28 dpLD: p<0.001; Figure 4E), indicating preserved inner retinal function. Notably, this preservation occurred despite reduced rod input, suggesting that altered MG function may contribute to maintained inner retinal responsiveness.

Photopic ERGs were used to assess cone function. At 14 dLD, cKO mice showed reductions in photopic b-wave amplitudes (−32%) comparable to light-damaged wildtypes (Figures 4G, S3F). However, at 28 dpLD, cKO mice exhibited a significant functional preservation relative to light-damaged wildtypes (p=0.032), indicating a potential delay in cone degeneration at this stage.

Together, these data demonstrate that MG-specific Dicer1 depletion in the Rlbp-Cre-Dicer-cKO mouse results in functional preservation of rod, cones, and the inner retina that was sustained up to 28d following light damage. Notably, inner retinal function remained fully preserved, highlighting a strong protective effect potentially associated with altered MG activity.

### Glast-Cre-driven Dicer-cKO in MG results in transient structural and functional preservation

Because the Rlbp1-Cre–driven Dicer-cKO partially affects the RPE ^61^, which is itself impacted by light damage (see review ^93^), we next assessed whether the observed phenotype is also found in the Glast-Cre-driven Dicer-cKO model, which selectively targets MG while sparing the RPE ^61^. The same experimental design was applied (Figure 5A).

**Figure 5:**
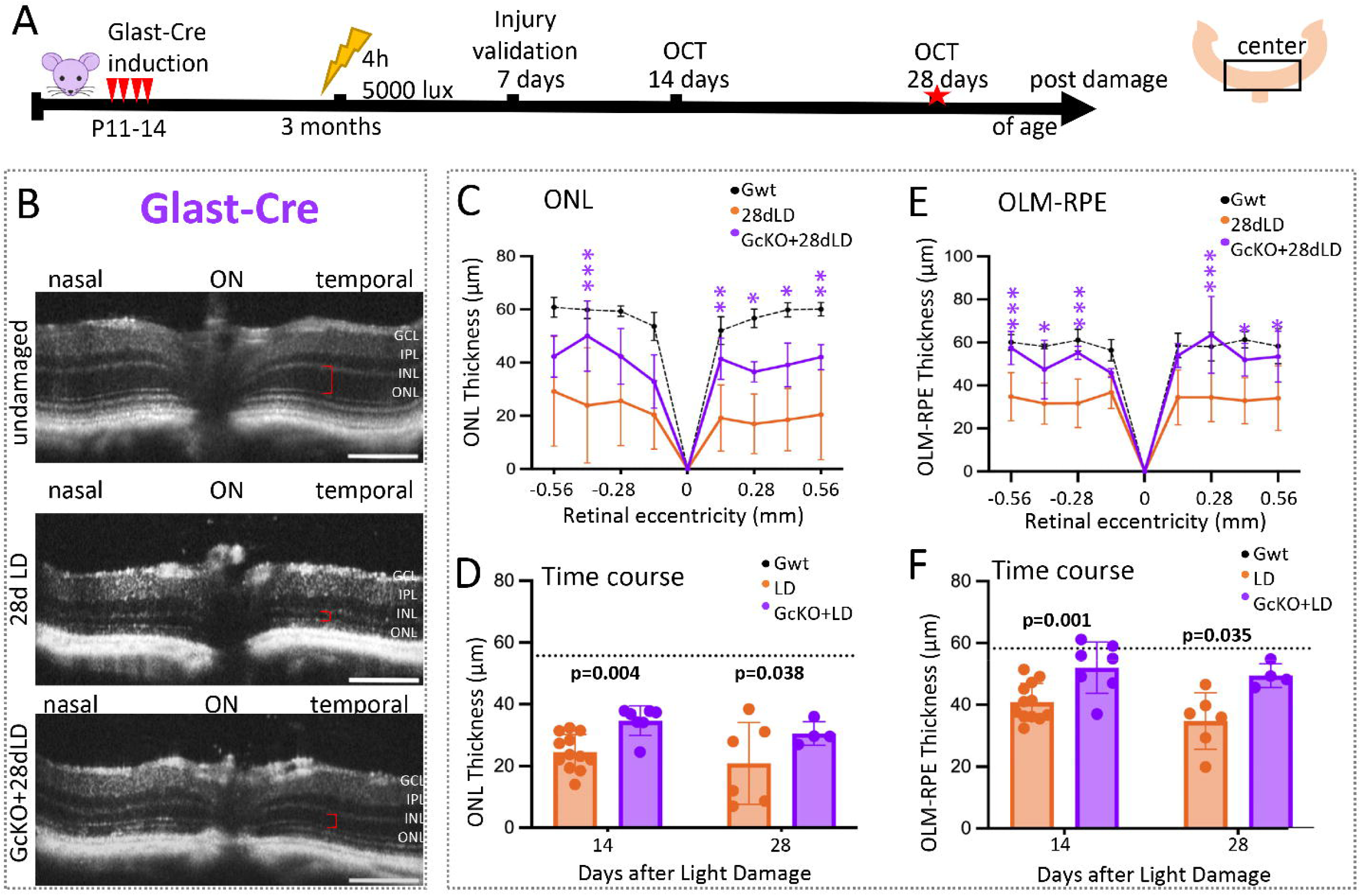
Light-damaged Glast-Cre: Dicer-cKO mice display a temporary preservation of retinal structure. A: Experimental design of the Optical Coherence Tomography (OCT) series. B: OCT images of center retinas (1300 μm diameter) at the nasal-temporal axis of undamaged Glast-Cre wildtype mice (Gwt), light-damaged Glast-Cre: Dicer-cKO (GcKO+LD), as well as light-damaged wildtypes (LD) 28 days post light damage (dLD). Red brackets indicate ONL thickness. **C-E:** Spider plots (C, E, nasal-temporal axis) and time course (D, F, averaged nasal-temporal and superior-inferior axis) of the thickness (diameter, μm) of the outer nuclear layer (ONL, C, D), and the outer limiting membrane/retinal pigment epithelium (OLM-RPE, E) of undamaged Glast-Cre wildtype mice (Gwt, n=6), light-damaged Glast-Cre: Dicer-cKO (GcKO+LD, n=4), as well as light-damaged wildtypes (LD, n=6) 14 and 28 days following light damage. Mean ± S.D.; significant differences between damaged wildtypes and cKOs are indicated using a two-way ANOVA with Tukey’s multiple comparisons correction; *: p≤0.05, **: p≤0.01 (C, E) and linear mixed-effects model with post hoc comparisons using estimated marginal means and Bonferroni correction for multiple comparisons in D, F; p-values for time-line graphs are given in the graphs. Undamaged wildtype values serve only as a reference/baseline. Scale bars in B: 200 µm. GCL: ganglion cell layer, IPL: inner plexiform layer, INL: inner nuclear layer, OPL: outer plexiform layer, ONL: outer nuclear layer, OLM: outer limiting membrane.

Similarly, as seen in the Rlbp-Cre: Dicer-cKOs, OCT imaging confirmed photoreceptor loss in Glast-Cre: Dicer-cKO retinas as early as 7d pLD, confirming effective damage induction (Figure S4). Moreover, also in the Glast-Cre driver Dicer-cKO, the retinal structure was preserved, particularly the outer retina (ONL and OLM-RPE, Figures 5B-F, S4).

We next assessed retinal function using ERG (Figure 6A). Scotopic a-wave amplitudes in undamaged Glast-Cre mice (∼235 μV) were comparable to those in Rlbp1-Cre controls. Moreover, similarly to the Rlbp-Cre strain, Glast-Cre: Dicer-cKO mice had significantly higher a-wave amplitudes 14 dpLD: (186 μV) compared to light-damaged wildtypes (p=0.001, Figures 6B, D, S5A), indicating preservation of rod function. However, this effect was not sustained at 28 dpLD (99 μV LD vs. 107 μV cKO, p=0.549; Figures 6C-D, S5A).

**Figure 6:**
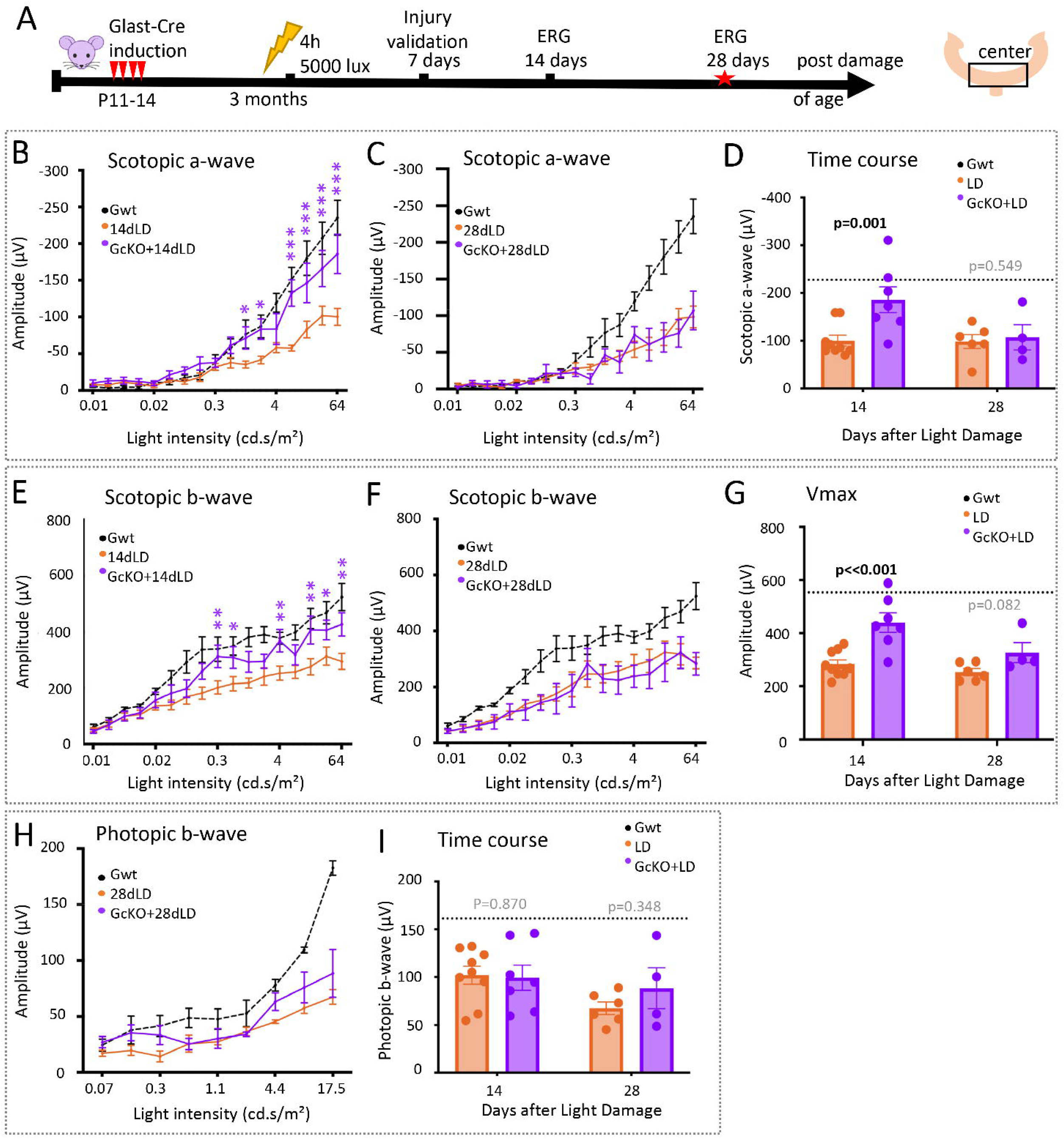
Light-damaged Glast-Cre: Dicer-cKOs display a temporary functional preservation. A: Experimental design of the electroretinography (ERG) series. B-I: Full-field ERG recordings showing scotopic a-wave amplitudes (B,C) and corresponding time line of the averaged amplitudes (D), scotopic b-wave amplitudes (E, F) and estimated saturated amplitudes (Vmax, responsiveness) using the Naka-Rushton equation (G), as well as photopic b-wave amplitudes (H) and corresponding time line of the averaged amplitudes (I) of undamaged Glast-Cre wildtypes (Gwt, n=5), light-damaged Glast-Cre: Dicer-cKOs (GcKO+LD) 14d (n=7), or 28dLD (n=4) as well as light damaged wildtypes (LD) 14d (n=9), or 28dLD (n=6). Mean ± S.E.M. Significant differences between damaged wildtypes and cKOs are indicated, using a two-way ANOVA with Tukey’s multiple comparisons correction; *: p≤0.05, **: p≤0.01, ***: p≤0.001 (B, C, E, F, H) and linear mixed-effects model with post hoc comparisons using estimated marginal means and Bonferroni correction for multiple comparisons in D, G, I; p-values for time-line graphs are given in the graphs. Undamaged wildtype values serve only as a reference /baseline.

Due to the alteration in input, the scotopic b-wave amplitudes (rod bipolar and MG function) were preserved 14d pLD in Glast-Cre: Dicer-cKO mice but not sustained 28d pLD (Figures 6E-G, S5B). The analysis of Vmax revealed a transient preservation of inner retinal responsiveness at 14 dLD, which was comparable to undamaged levels and very similar to the outcome seen in the Rlbp-Cre-Dicer-cKO (Figures 6G, S5C). This inner retinal preservation in the Glast-Cre model is, however, more restricted and only temporary.

Photopic ERG recordings showed no significant preservation of cone function in Glast-Cre: Dicer-cKO mice at any time point (Figures 6H-I, S6), although a trend toward improved function was observed, similar to that seen in the Rlbp1-Cre model.

Together, these results indicate that MG-specific Dicer1 deletion using the Glast-Cre driver confers transient preservation of rod and inner retinal function following light damage, primarily within the first two weeks. In contrast to the Rlbp1-Cre model, this protective effect is not sustained, although the retinal structure was preserved.

### Histology confirms reduced structural damage to rod photoreceptors

To further evaluate retinal structure, we performed histological analyses of undamaged and light-damaged retinas 28 days post-light damage (dLD). Given the inherent variability of the light damage model, lesion localization did not always precisely match the central regions assessed by OCT. Nevertheless, consistent with *in vivo* findings, light-damaged cKO retinas exhibited less extensive structural damage compared to light-damaged wildtypes (Figure 7A. showing representative bright field images of whole retinal cross sections).

**Figure 7:**
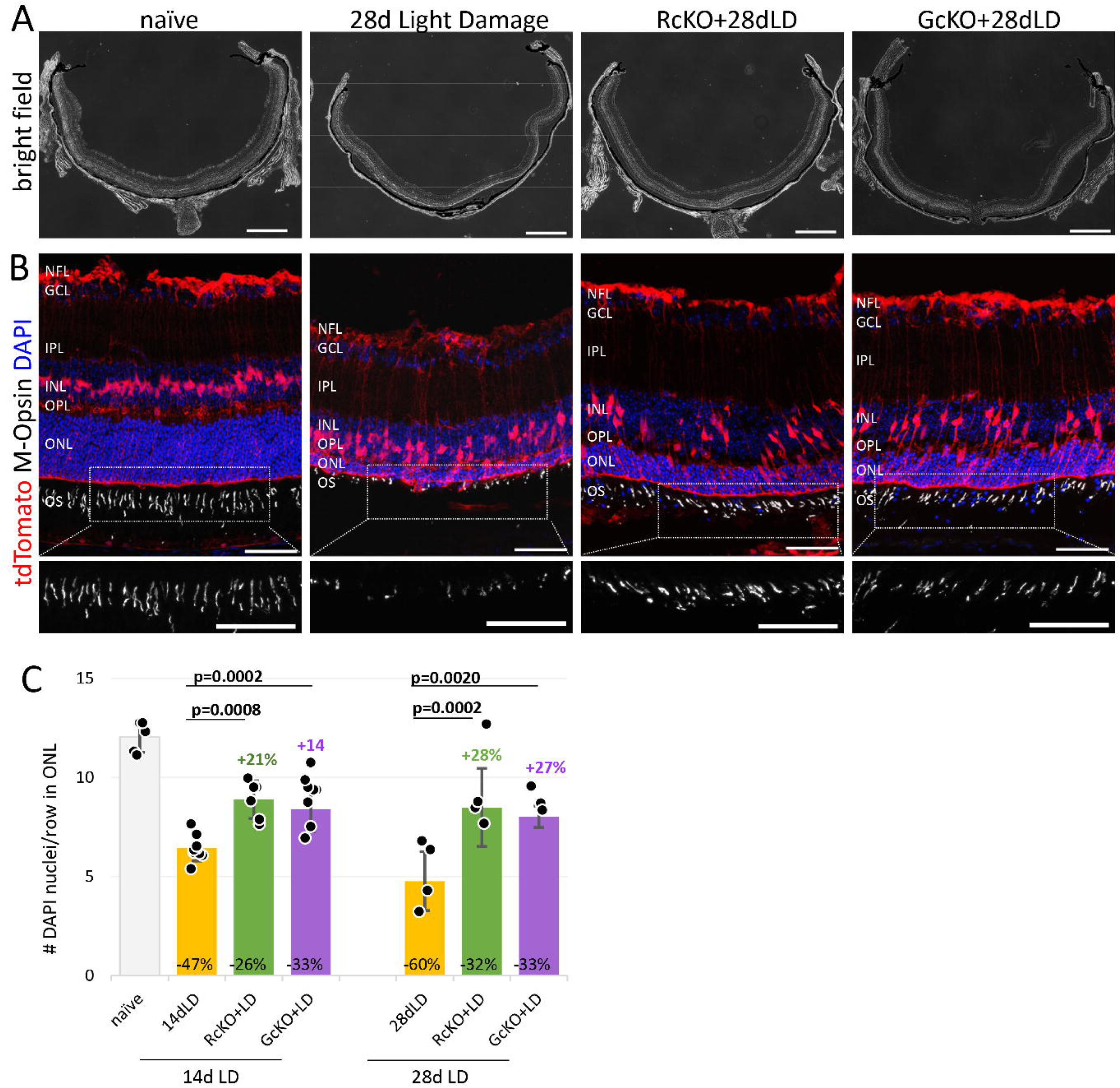
Light-damaged Dicer-cKO mice retain rods and cones. A: Bright field images of whole retinal cross sections of an undamaged (naïve) mouse, as well as a light-damaged wildtype (28d Light Damage), Rlbp-Cre: Dicer-cKO (RcKO+28dLD), and Glast-Cre: Dicer-cKO mouse (GcKO+28dLD), 28 days after damage. B: Immunofluorescent labeling with antibodies against M-Opsin of DAPI counterstained representative damage areas with endogenously labeled tdTomato+ MG of an undamaged (naïve) wildtype, as well as a light-damaged wildtype (28d Light Damage), Rlbp-Cre: Dicer-cKO (RcKO+28dLD), and Glast-Cre: Dicer-cKO mouse (GcKO+28dLD) 28 days after damage. The insets show magnified areas with cone outer segments. C: Number of DAPI+ nuclei per row in the ONL of damaged areas in undamaged mice (naïve, n=5, reference/baseline) as well as damaged wildtypes (LD, 14d: n=9; 28d: n=4) and cKOs (Rlbp-Cre [RcKO] 14d: n=6, 28d: n=4; and Glast-Cre [GcKO] 14d: n=8, 28d: n=3), 14 and 28 days after light damage (14dLD, 28d LD, respectively). Mean ± S.D. Two-way ANOVA with Dunnett’s multiple comparisons correction comparing each condition to light-damaged wildtype within each time point. Undamaged naïve data serves solely as reference. Scale bars in A: 500 μm, in B 50 µm. NFL: Nerve Fiber Layer, GCL: ganglion cell layer, IPL: inner plexiform layer, INL: inner nuclear layer, OPL: outer plexiform layer, ONL: outer nuclear layer, OS: outer segments.

To assess photoreceptor survival, sections were labeled with DAPI and M-opsin (Figure 7B). As the majority of DAPI+ nuclei in the ONL represent rods ^94^, nuclei counts per row were used as an estimate of rod numbers. Quantification revealed significantly higher numbers of DAPI+ nuclei in both cKO strains compared to light-damaged wildtypes at 14 dLD and 28 dLD (Figure 7C), indicating reduced rod loss.

Cone integrity was evaluated using M-opsin labeling. M-Opsin antibodies label medium-wavelength sensitive opsin, which is evenly distributed in the mouse retina and allows assessment of cone health. In undamaged retinas, M-opsin+ outer segments appeared long and well-organized. Light-damaged wildtypes displayed markedly shortened or absent segments (Figure 7B, insets), in accordance with previous reports ^95^. In contrast, light-damaged cKO retinas retained numerous M-opsin+ segments, although somewhat reduced and shortened. No clear differences were observed between the two cKO strains.

Together, these histological data corroborate the *in vivo* OCT findings and indicate that MG-specific Dicer1 deletion results in reduced photoreceptor alterations. Importantly, both MG-targeting strategies (Rlbp-Cre and Glast-Cre driven MG-specific Dicer-cKO) yielded comparable structural outcomes.

### MG Dicer1 loss drives functional preservation independent of developmental timing and pre-existing retinal state

To determine whether the observed phenotype is independent of the timing and context of MG manipulation, we analyzed a third MG-targeting model, the Ascl1-Cre: Dicer-cKO. Depending on the time point of Cre recombination, this line can target all MG as well as some late-born neuron populations ^96, 97^. We originally developed this line to study Dicer1 function in postnatal retinal progenitor cells (RPCs). Early postnatal Dicer loss resulted in an altered composition of the postnatal retina, but more importantly, in delayed MG maturation and an unusually slow neuronal degeneration in the adult mouse ^57^. This suggested that the Dicer1 depletion in the glial progenitors/precursors had an impact on the glial state.

To assess whether a similar MG-driven mechanism may operate in this model, Ascl1-Cre Dicer-cKO mice were subjected to light damage at around 1.5 months of age following Cre induction at P1–3 (Figure 8A). In the adult retina, tdTomato+ Sox2+ MG are found throughout the retina in wildtype as well as cKO retinas. Their nuclei are in the INL, and their endfeet form the OLM (Figure 8B, white arrows and arrowheads, respectively).

**Figure 8:**
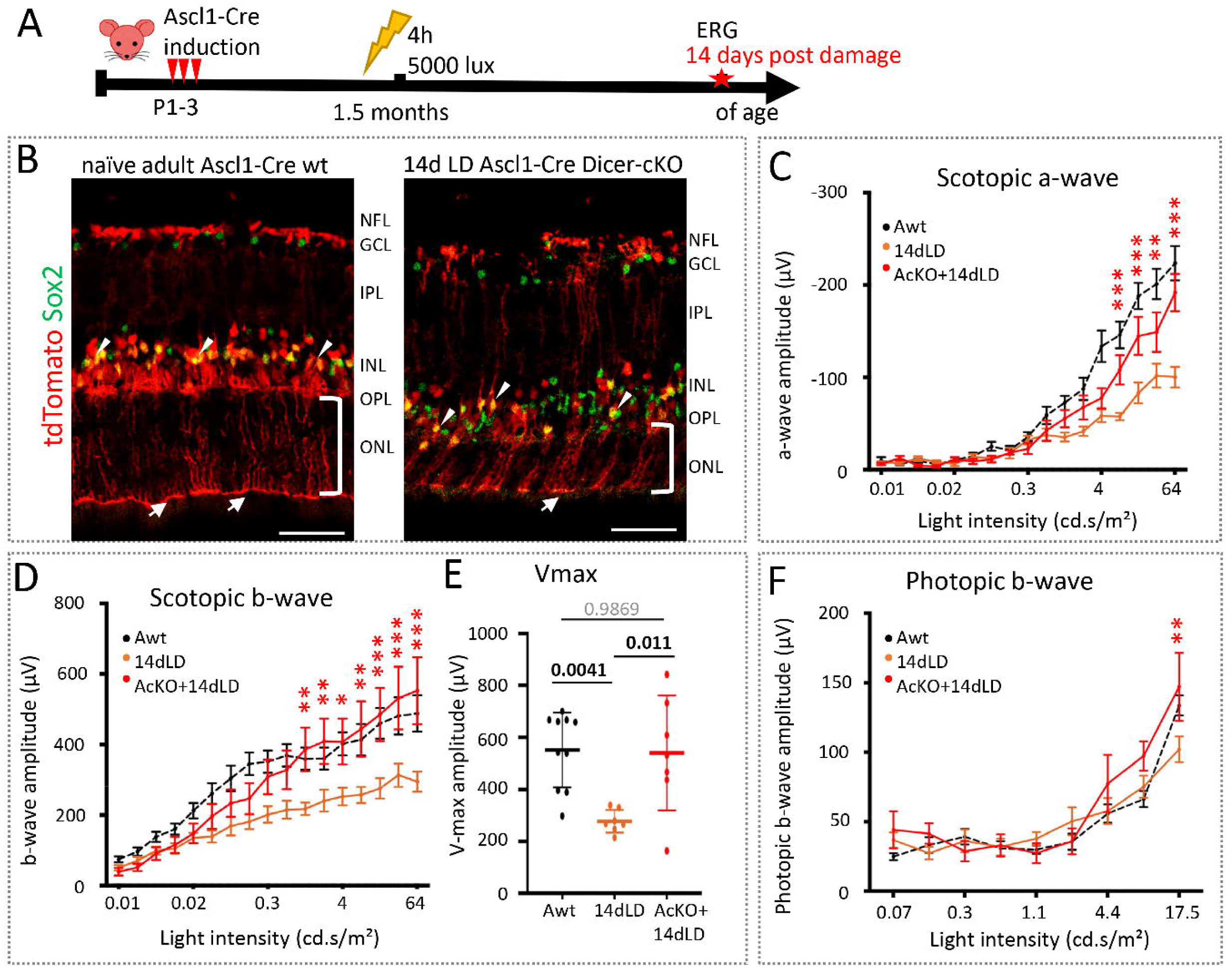
Light-damaged Ascl1-Cre: Dicer-cKO mice display functional preservation. A: Experimental design of the electroretinography (ERG) study. B: Immunofluorescent labeling with antibodies against Sox2 of retinal cross sections of representative damage areas with endogenously labeled tdTomato+ MG in an undamaged naïve Ascl1-Cre wt mouse and an Ascl1-Cre-Dicer-cKO 14d pLD. Arrowheads indicate GFAP+ MG in the INL; arrows indicate the ONL formed by MG endfeet; brackets indicate ONL thickness. C-F Full-field ERG recordings showing scotopic a-wave amplitudes (C), scotopic b-wave amplitudes (D), and estimated saturated amplitudes (Vmax, responsiveness) using the Naka-Rushton equation (E), as well as photopic b-wave amplitudes (F) of Ascl1-Cre wildtype (Awt, n=10), light-damaged wildtypes (14dLD, n=7), and light-damaged Ascl1-Cre-Dicer-cKO mice (AcKO+14dLD, n=7) 14 days after damage. Mean ± S.E.M. Significant differences between damaged wildtypes and cKOs are indicated using a two-way ANOVA with Tukey’s multiple comparisons correction; *: p≤0.05, **: p≤0.01, ***: p≤0.001 in C, D, F, and one-way ANOVA with Tukey’s multiple comparisons correction (E, p-values given in the graph). Undamaged wildtype values serve only as a reference /baseline. Scale bars: 50 µm. NFL: Nerve Fiber Layer, GCL: ganglion cell layer, IPL: inner plexiform layer, INL: inner nuclear layer, OPL: outer plexiform layer, ONL: outer nuclear layer, OLM: outer limiting membrane.

Our previous study demonstrated that at this age, although no overt structural abnormalities were present, Ascl1-CreER Dicer1-cKO retinas already exhibited reduced rod and inner retinal function (approximately 30% reductions in scotopic a- and b-wave amplitudes) resulting from developmental alterations, whereas cone function remained preserved ^57^. Despite this baseline functional deficit, the mice were subjected to light damage to determine whether MG-specific Dicer1 deletion influenced the retinal response to injury in the presence of pre-existing dysfunction. Following light damage, cKO retinas exhibited thinning of the outer nuclear layer (ONL), confirming successful injury induction (Figure 8B, brackets).

Functional analyses were performed 14 days post-light damage and compared to age-matched light-damaged wildtypes (Figures 8C, S1C-D). Scotopic a-wave amplitudes were significantly higher in cKO mice compared to light-damaged wildtypes (14 dpLD: 191 μV vs. 100 μV, p<0.0001), indicating partial preservation of rod function. Similarly, scotopic b-wave amplitudes were significantly increased in cKO mice (552 μV vs. 294 μV, p<0.0001; Figure 8D), reflecting a preserved inner retinal function.

Consistent with these findings, Vmax values derived from Naka-Rushton fits were significantly higher in cKO mice compared to light-damaged wildtypes p=0.0110; Figure 8E), confirming preserved inner retinal responsiveness. Also, photopic ERGs showed a preserved cone function in cKO mice (147 μV vs. 102 μV, p=0.0010; Figure 8F).

Together, these data demonstrate that MG-specific Dicer1 deletion is consistently associated with preservation of retinal function, particularly within the inner retina, even in a developmentally altered retinal context. Across all three Cre driver lines, Dicer1 manipulation in MG represents the common experimental feature underlying this phenotype. Importantly, the protective response was observed despite differences in the developmental stage, timing of Dicer1 deletion, and baseline retinal function, highlighting the robustness of this association. Notably, the absence of further functional decline following light damage suggests that Dicer1-deficient MG respond differently to retinal injury than their wild-type counterparts, although the molecular basis of this altered response remains to be determined.

### Prior injury and light preconditioning do not reproduce the Dicer1-dependent protective phenotype

Given the stable functional phenotype observed in Ascl1-Cre: Dicer-cKO mice after light damage ^57^, together with the presence of baseline retinal dysfunction in all MG-specific Dicer1-cKO models prior to injury ^61^, we asked whether pre-existing retinal dysfunction or prior injury exposure could account for the observed neuroprotective phenotype.

To address this question, we employed two complementary experimental paradigms. First, we established a double light damage paradigm to reproduce the degree of functional impairment observed in the Ascl1-Cre; Dicer1-cKO model (approximately 30% reduction in rod responses with mild cone dysfunction). Mice received two identical light damage exposures (5,000 lux for 4 hours) separated by a 3-day interval (Figure 9A). This interval was selected because rod dysfunction is already present at this stage, whereas retinal structure remains largely preserved (Figures S1E–G). Retinal function was assessed by ERG 7 days after the initial exposure (4 days after the second light damage) and compared with mice subjected to a single light damage.

**Figure 9:**
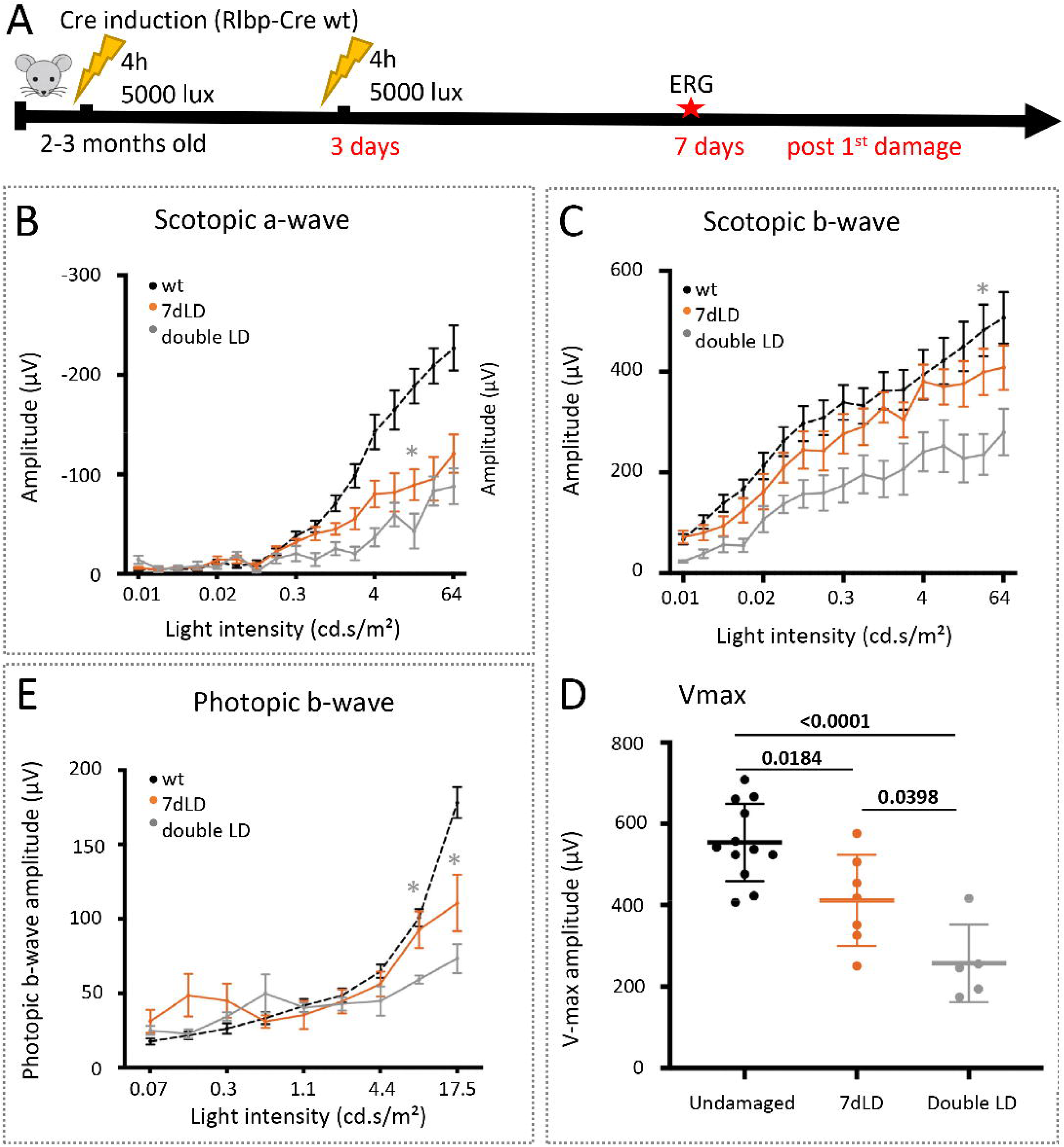
Sequential damage does not result in functional preservation. A: Experimental design of the electroretinography (ERG) study. B-E: Full-field ERG recordings showing scotopic a-wave amplitudes (B), scotopic b-wave amplitudes (C), and estimated saturated amplitudes (Vmax, responsiveness) using the Naka-Rushton equation (D), as well as photopic b-wave amplitudes (E) of Rlbp-Cre wildtype (Rwt, n=11), light-damaged wildtypes (single damage, 7dLD, n=6), and wildtype mice light-damaged a second time, 3 days after the first damage (double LD 7d, n=5). Mean ± S.E.M. Significant differences between single and double-damaged wildtypes are indicated using a two-way ANOVA with Tukey’s multiple comparisons correction; *: p≤0.05, **: p≤0.01, ***: p≤0.001 in B, C, E, and one-way ANOVA with Tukey’s multiple comparisons correction (D, p-values given in the graph). Undamaged wildtype values serve only as a reference /baseline.

Scotopic a-wave amplitudes in double-damaged mice were comparable to those in single-damaged mice (−88 μV vs. −120 μV, p=0.20642; Figure 9B), with differences primarily at lower intensities. Scotopic b-wave amplitudes were likewise similar between groups (double LD: ∼280 μV vs. single LD: ∼407 μV, p=0.1087; Figure 9C). Consistent with these findings, Vmax was not preserved and instead showed a greater reduction following double light damage than after a single exposure (p=0.0398; Figure 9D). Photopic b-wave amplitudes were similarly reduced in both groups (73 μV vs. 110 μV, p=0.0172; Figure 9E). Thus, prior injury exposure did not preserve rod, cone, or inner retinal function.

Second, we evaluated a classical light preconditioning paradigm consisting of a mild light exposure (1,000 lux for 2 hours) followed 3 days later by the standard light damage paradigm (5,000 lux for 4 hours) (Figure 10A). Functional assessment again revealed no evidence of protection. Scotopic a- and b-wave amplitudes, Vmax, and photopic responses were comparable between preconditioned and non-preconditioned mice (Vmax, p = 0.9172; Figures 10B–E).

**Figure 10:**
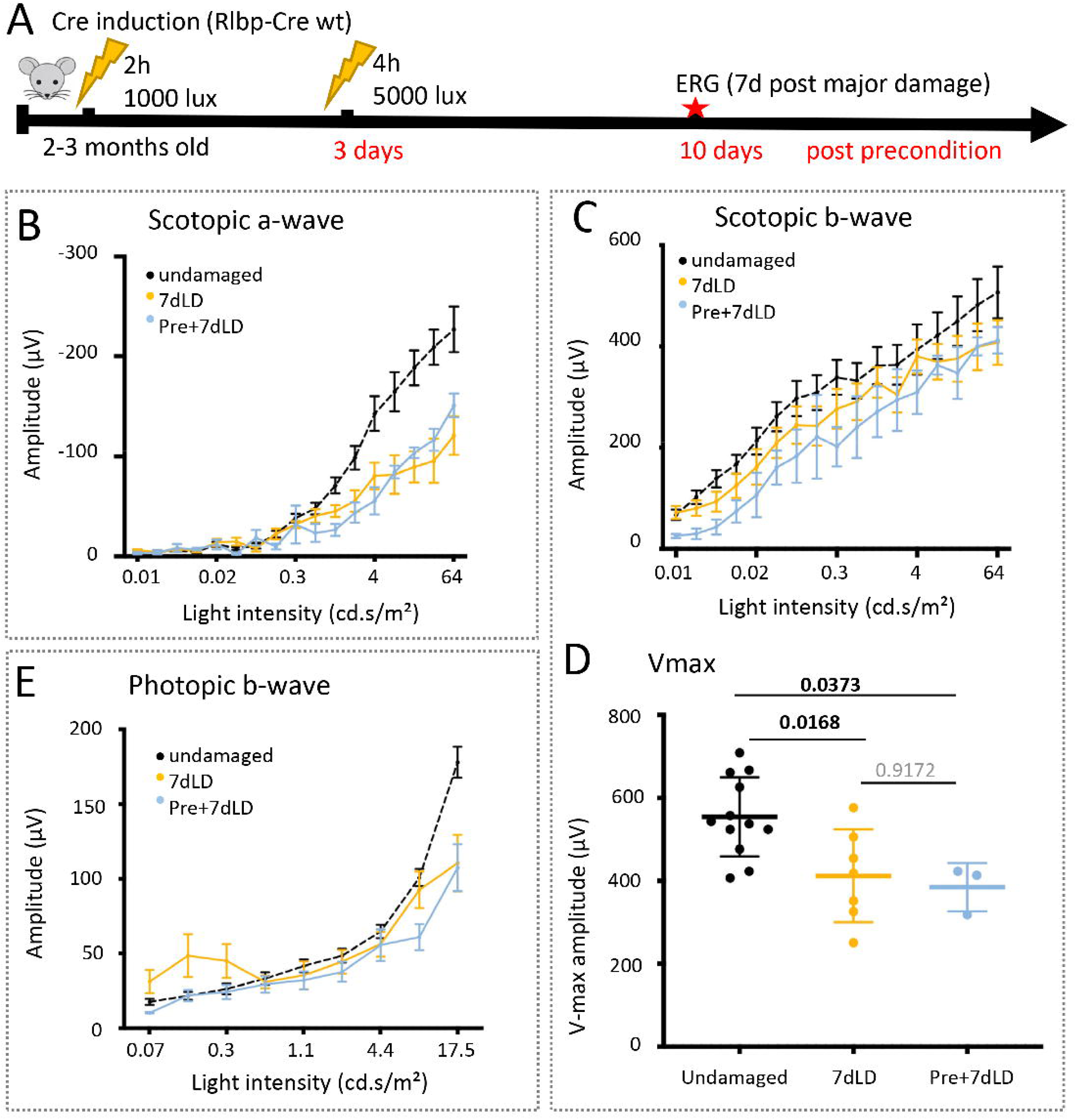
Preconditioning does not result in functional preservation. A: Experimental design of the electroretinography (ERG) study. B-E: Full-field ERG recordings showing scotopic a-wave amplitudes (B), scotopic b-wave amplitudes (C) and estimated saturated amplitudes (Vmax, responsiveness) using the Naka-Rushton equation (D), as well as photopic b-wave amplitudes (E) of Rlbp-Cre wildtype (Rwt, n=11), light-damaged wildtypes (single damage, 7dLD, n=6), and wildtype mice which were preconditioned 3 days before light-damage (pre+7dLD, n=3). Mean ± S.E.M. Significant differences between damaged and preconditioned wildtypes were evaluated using a two-way ANOVA with Tukey’s multiple comparisons correction in B, C, and E, and a one-way ANOVA with Tukey’s multiple comparisons correction (D; p-values given in the graph). Undamaged wildtype values serve only as a reference /baseline.

Together, neither repeated light damage nor classical light preconditioning reproduced the functional preservation observed in MG-specific Dicer1-cKO mice. In particular, inner retinal function was not maintained under either condition, indicating that the protective phenotype is unlikely to be explained by prior injury exposure or preconditioning and instead is associated with MG-specific Dicer1 deletion.

### Dicer1-deficient and preconditioned retinas exhibit reduced GFAP immunoreactivity after light damage

To evaluate the Müller glial response following light damage, we performed GFAP immunostaining in undamaged, light-damaged, and MG-specific Dicer1-cKO retinas. In light-damaged wildtype retinas, GFAP immunoreactivity was markedly increased and extended from the inner retina into Müller glial processes, consistent with the induction of reactive gliosis (Figures 11A–D and S6).

**Figure 11:**
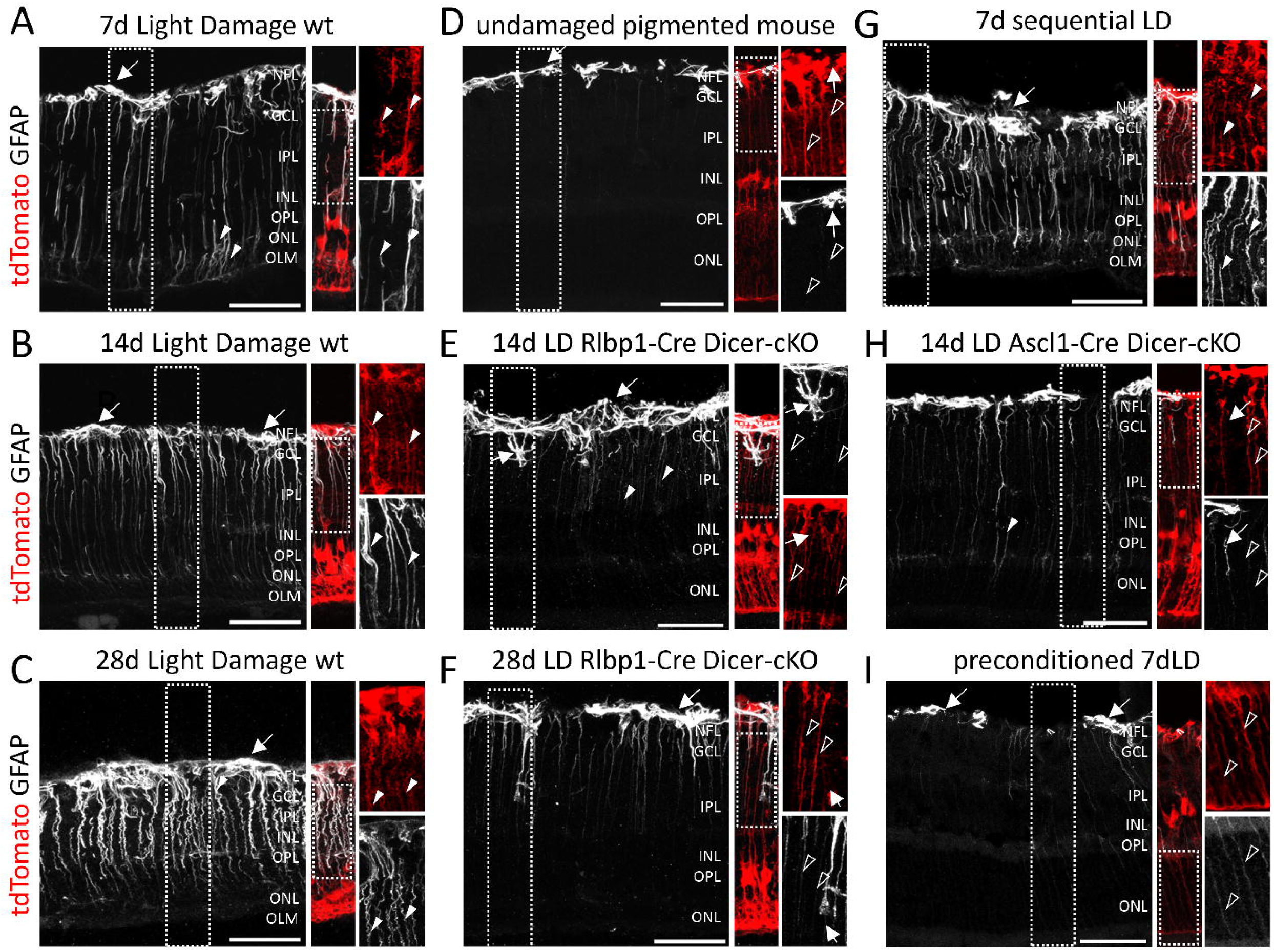
Müller glia of light-damaged Dicer-cKO mice lack GFAP immunoreactivity. A-I: Immunofluorescent labeling with antibodies against GFAP of retinal cross sections of representative damage areas with endogenously labeled tdTomato+ MG of a wildtype mouse 7 days (A), 14 days (B), or 28 days (C) after light damage, of an undamaged pigmented mouse (D), of a Rlbp-Cre: Dicer-cKO mouse 14days after light damage (E) or 28 days after light damage (F); of a wildtype mouse which underwent sequential damage (3d+4d, G), of a Ascl1-Cre: Dicer-cKO mouse 14 days after light damage (H) as well as of a preconditioned wildtype mouse 7 days after light damage (I). Filled arrowheads indicate GFAP+ MG processes, unfilled arrowheads indicate MG processes with very low or no GFAP co-localization, and arrows indicate tdTomato-negative astrocytes. Scale bars: 50 µm. NFL: Nerve Fiber Layer, GCL: ganglion cell layer, IPL: inner plexiform layer, INL: inner nuclear layer, OPL: outer plexiform layer, ONL: outer nuclear layer, OLM: outer limiting membrane.

In contrast, MG in light-damaged Dicer1-cKO retinas exhibited markedly reduced GFAP immunoreactivity. GFAP expression remained largely confined to the GCL/NFL, with minimal extension into radial MG processes, consistent with a reduced GFAP-associated injury response. This phenotype was consistently observed across all MG-specific Cre lines despite differences in the extent of structural and functional preservation (Figures 11E–F, H and S7). In some cKO retinas, prominent GFAP-positive, tdTomato-negative astrocytic processes were observed (arrows, Figures 11E–F, H), clearly distinguishable from MG by the reporter, indicating that astrocytes remained reactive despite the reduced GFAP immunoreactivity in MG.

Interestingly, the double light damage paradigm, which was designed to reproduce the degree of retinal dysfunction present in the Ascl1-Cre; Dicer1-cKO model at the time of injury, resulted in robust GFAP immunoreactivity throughout Müller glial processes (Figure 11G). Thus, pre-existing retinal dysfunction alone was insufficient to attenuate GFAP expression following light damage. In contrast, retinas subjected to the low-intensity light preconditioning paradigm displayed reduced GFAP immunoreactivity (Figure 11I), consistent with previous reports.98 Despite this reduction, preconditioned retinas did not exhibit preservation of retinal function.

Together, these findings demonstrate that MG-specific Dicer1 deletion is consistently associated with reduced GFAP immunoreactivity following light damage. However, because reduced GFAP expression was also observed after light preconditioning without accompanying functional preservation, diminished GFAP immunoreactivity alone is insufficient to explain the neuroprotective phenotype observed in Dicer1-cKO retinas.

Overall, MG-specific Dicer1 deletion was associated with structural preservation and sustained maintenance of inner retinal function following light damage across three independent MG-targeting models. Although the magnitude and duration of protection differed among the Cre lines, neither prior injury exposure nor light preconditioning reproduced the functional phenotype. A schematic overview summarizing the experimental paradigms and principal findings is provided in Figure 12.

**Figure 12:**
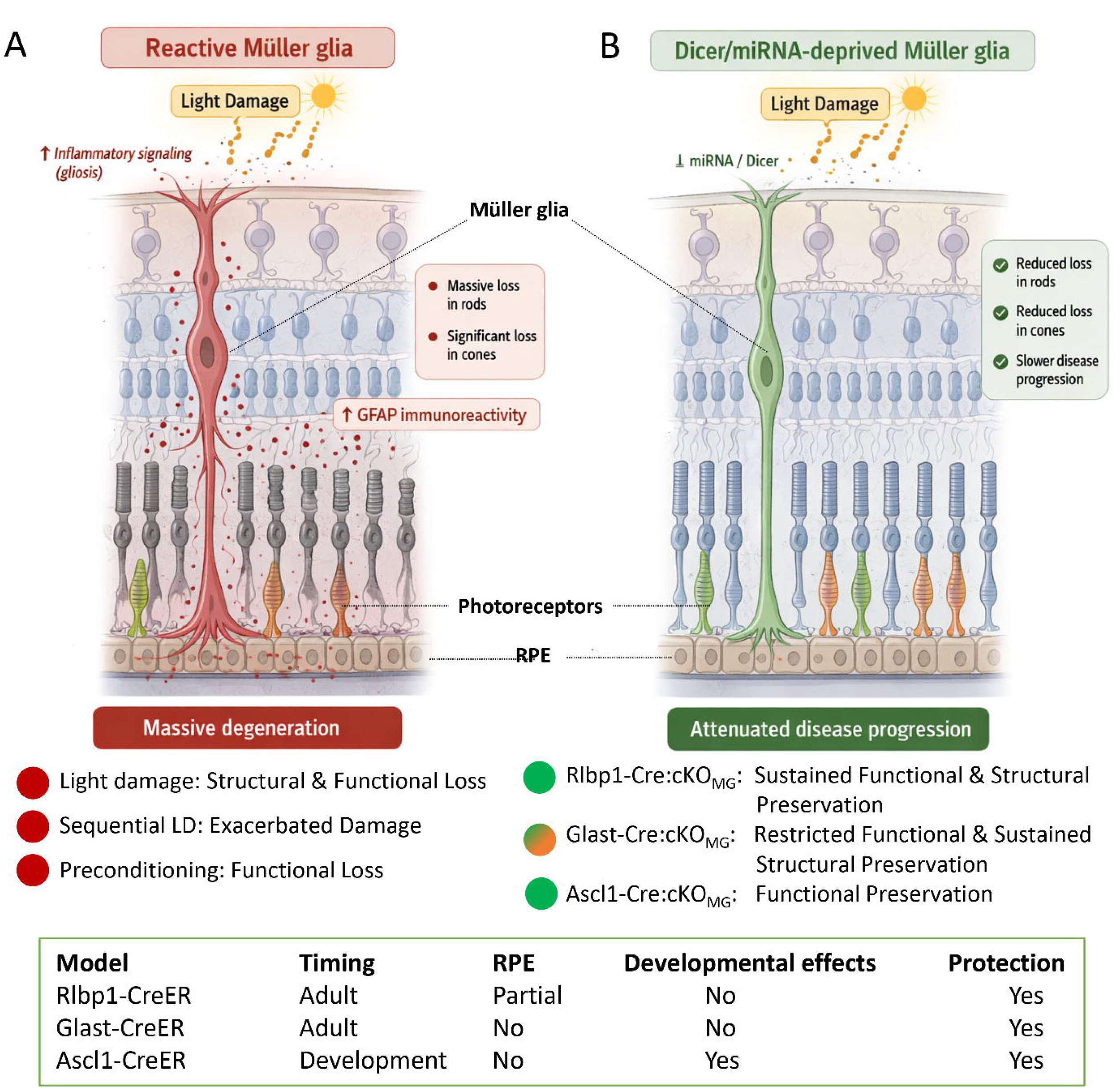
Major findings of light-damaged wildtypes and MG-specific cKOs at a glance. A: Simplified illustration of major events in degenerating retinas after light damage, showing dying photoreceptors and reactive (GFAP+) MG. B: Simplified illustration of events predominantly found in MG-specific Dicer1-cKO retinas after light damage, showing preserved photoreceptors and non-reactive MG. C: Overview of mouse strains and damage paradigms used, categorized by and mapped to the major phenotypes shown in A and B. Red dots symbolize loss of structure and function; green dots preservation, and orange coloring restricted preservation. A and B were generated using OpenAI and do not represent correct retinal circuits.

## Discussion

The role of Müller glia (MG) in retinal disease progression has been investigated for decades. Similar to astrocytes throughout the central nervous system, MG undergo reactive gliosis following injury and contribute to inflammation and secondary neuronal degeneration ^98–102^. However, the molecular mechanisms that determine how MG respond to injury and influence disease progression remain incompletely understood.

In the present study, we demonstrate that MG-specific Dicer1 deletion is associated with structural and, more importantly, functional preservation following light-induced retinal injury. This phenotype was reproducible across three independent MG-targeting Cre driver lines despite differences in the developmental stage and timing of Dicer1 deletion. Structural preservation was observed in both the Rlbp1-CreER- and Glast-CreER-driven models, whereas sustained functional preservation was most pronounced in the Rlbp1-CreER model. Importantly, neither repeated light damage nor light preconditioning reproduced this phenotype, indicating that the observed protection is linked to Dicer1-deficient MG rather than prior retinal injury.

Together, these findings support an active role for Müller glia in shaping the retinal response to injury and identify Dicer1 as an important determinant of this response.

### A moderate light damage model for pigmented transgenic mice

Light-induced retinal degeneration is a well-established experimental model for studying photoreceptor loss and reproducing important features of retinitis pigmentosa (RP) and age-related macular degeneration (AMD) ^3–5, 7, 8, 93^. Historically, most studies have relied on albino animals due to their high susceptibility to light-induced injury ^20, 35, 103–106^. However, albino retinas differ from pigmented retinas in several important respects, including retinal development, visual acuity, and baseline GFAP expression, suggesting a pre-stressed retinal environment ^36–38^. These differences raise concerns regarding the physiological relevance of albino models for studying human retinal disease. Conversely, most transgenic mouse lines, including the Dicer1^fl/fl^ strain used in this study, are maintained on the C57BL/6 background, which is relatively resistant to light damage because of the protective RPE65^Met450^ variant ^107, 108^. Consequently, reliable injury induction requires specialized light damage paradigms.

To overcome these limitations, we established a moderate light damage paradigm in pigmented transgenic mice carrying the RPE65^Leu450^ variant, enabling inducible MG-specific Dicer1 deletion together with longitudinal structural and functional assessment. To our knowledge, only a limited number of studies have employed light damage paradigms in pigmented S129 mice, and these generally used substantially higher light intensities (>10,000 lux) that produced more severe acute injury ^80, 95, 107^.

The light damage paradigm used in this study recapitulates the temporal progression of degeneration observed in human retinal diseases, with early rod dysfunction followed by secondary cone degeneration and delayed impairment of the inner retina. This progression resembles that seen in AMD and RP ^25–27, 42, 99, 109–112^, supporting the concept that non-cell-autonomous mechanisms make important contributions to retinal degeneration.

Notably, we found that functional decline preceded detectable structural loss and that inner retinal dysfunction emerged earlier than previously recognized. This observation was uncovered through Vmax analysis, which is not routinely included in studies of retinal degeneration but provides additional insight into inner retinal function that is not captured by conventional ERG parameters alone. Our findings therefore highlight the value of incorporating Vmax measurements, as structural and molecular changes do not necessarily reflect functional integrity.

Overall, this moderate light damage paradigm provides a physiologically relevant platform for investigating retinal degeneration in pigmented transgenic mice while enabling selective interrogation of MG contributions to disease progression.

### Müller glia shape the retinal response to injury: Dicer1 deletion reveals a neuroprotective phenotype

One of the principal findings of this study is that altering Dicer1 function in MG substantially changes the retinal response to injury. MG-specific Dicer1 deletion was associated with preservation of rod and cone function, maintenance of inner retinal responses, and attenuation of secondary degeneration following injury. Importantly, protective effects were consistently observed across three independent MG-targeting Cre driver lines, despite differences in the magnitude and duration of preservation. Interestingly, even in the Ascl1-Cre model, in which developmental Dicer1 deletion produces baseline retinal abnormalities and impaired visual function ^57^, light damage did not further exacerbate the phenotype. This observation is consistent with our previous findings that, despite these developmental defects, retinal degeneration in this model progressed unexpectedly slowly ^57^.

Together, these observations suggest that Dicer1-deficient MG respond differently to retinal injury, thereby limiting further functional decline. Notably, preservation of inner retinal function was the most consistent finding across all three models, suggesting that altered MG responses preferentially preserve retinal circuitry despite ongoing photoreceptor degeneration. Although the presence of immature or partially differentiated MG populations in this model may contribute to the observed phenotype^57^, further studies will be required to determine the underlying mechanisms.

### Dicer1 deletion modifies the Müller glial injury response

In our previous characterization of the MG-specific Rlbp1-CreER; Dicer1 cKO mouse, we found that GFAP expression was not induced, even under conditions of advanced retinal degeneration ^60^. Because GFAP is a well-established marker of reactive gliosis and has been associated with inflammation, glial hypertrophy, scar formation, and secondary neuronal degeneration ^24, 44, 46, 113–116^, these findings suggested that Dicer1 depletion fundamentally alters the canonical MG injury response to retinal injury. Nevertheless, GFAP represents only one aspect of reactive gliosis, and the molecular mechanisms linking Dicer1 loss to the observed neuroprotective phenotype remain unknown.

The concept of secondary degeneration provides an important framework for interpreting these findings. While the initial phase of light-induced retinal injury is driven primarily by direct phototoxicity and oxidative stress ^5, 15^, subsequent neuronal loss is thought to involve non-cell-autonomous mechanisms, including inflammatory signaling and glia-neuron interactions ^117, 118^. Our findings are consistent with the possibility that Dicer1-deficient MG modify this secondary phase of degeneration. Notably, reduced GFAP expression alone is unlikely to account for the observed protection, as light preconditioning similarly decreased GFAP immunoreactivity without preserving retinal function. Together, these findings indicate that Dicer1 depletion alters aspects of the MG injury response beyond GFAP expression alone.

Although the molecular basis of this altered injury response remains unresolved, several mechanisms merit further investigation. Müller glia regulate numerous processes essential for retinal homeostasis, including cytokine signaling, metabolic support, ion buffering, glutamate homeostasis, and intercellular communication ^18, 40–43, 119^, all of which could influence neuronal survival following injury. Among the candidate pathways, JAK/STAT3 signaling is particularly interesting because it is a well-established regulator of reactive gliosis in both the CNS and retina.^91, 120–128^. Future studies combining transcriptomic analyses with functional validation will be required to define how Dicer1 regulates the Müller glial injury response and promotes retinal preservation.

### Müller glia as the primary driver of the protective phenotype

Although the Rlbp1-CreER line also exhibits partial recombination in the RPE, the protective phenotype was observed across all three MG-targeting Dicer1-cKO models, including lines without detectable RPE involvement. Because these models differ in recombination pattern, developmental timing, and baseline retinal phenotype, the shared response is most consistently explained by Dicer1 deletion in MG. Nevertheless, we cannot exclude the possibility that partial Dicer1 deletion in the RPE contributes to the greater durability of structural and functional preservation observed in the Rlbp1-CreER model. Resolving this possibility will require RPE-specific Dicer1 manipulation and direct assessment of RPE function.

## Conclusion

In summary, we established a moderate light damage paradigm for pigmented transgenic mice that enables detailed investigation of MG contributions to retinal degeneration. Using this model, we show that MG-specific Dicer1 deletion is associated with structural preservation and, more importantly, sustained maintenance of retinal function following injury, supporting an active role for MG in shaping disease progression.

Notably, maintenance of inner retinal function despite ongoing photoreceptor degeneration suggests that Dicer1-deficient Müller glia influence not only neuronal survival but also the integrity of retinal circuitry. This is of high relevance as maintenance of downstream retinal circuitry is critical for emerging restorative approaches, including gene augmentation, optogenetics, cell replacement therapies, and retinal prosthetics. Although the mechanisms underlying this phenotype remain to be defined, the altered MG response observed following Dicer1 deletion provides a foundation for future studies aimed at identifying the molecular pathways that promote retinal resilience.

Collectively, our findings identify Dicer1 as an important regulator of the Müller glial injury response and establish Müller glia as active determinants of retinal resilience following injury.

## Limitations and Future Directions

Several limitations should be considered when interpreting the present findings. First, this study primarily provides a functional and phenotypic characterization of MG-specific Dicer1 deletion following retinal injury. Although Dicer1 depletion was consistently associated with structural and functional preservation, the molecular mechanisms underlying this phenotype remain unresolved. In particular, the downstream signaling pathways and Dicer1-dependent regulatory mechanisms responsible for the altered MG injury response were not investigated and warrant future study.

Second, while the three independent MG-targeting mouse lines provide strong evidence that the observed phenotype is primarily MG-driven, the individual models differ somewhat in their developmental timing and cellular specificity. Although these differences did not alter the overall direction of the phenotype, they may contribute to differences in the magnitude or duration of protection. In particular, partial recombination of the retinal pigment epithelium in the Rlbp1-CreER line could have influenced the sustained protective effects observed in this model and should be addressed using RPE-specific approaches.

Finally, although the light damage paradigm reproduces many features of retinal degeneration, it represents an acute injury model and therefore does not fully capture the complexity of chronic human retinal diseases. Future studies will be needed to determine whether similar Müller glial responses occur in inherited and age-related models of retinal degeneration.

An important next step will be to define the molecular mechanisms downstream of Dicer1 that mediate the observed phenotype. This includes identifying the relevant Dicer1-dependent small RNAs, their target pathways, and the signaling networks that regulate the transition between detrimental and protective MG states. Such studies will help determine whether selective modulation of these pathways can reproduce the beneficial effects observed following Dicer1 deletion while preserving normal MG function.

## Supporting information

Larbi 2026_Supplement revision

## Acknowledgements

The authors thank Dr. Suresh Viswanathan for advice on ERG-related questions and Monica Andrade for technical assistance with this study. This study was funded by the New York Empire Innovation Program Grant to S.G.W, the National Eye Institute (NEI, R01 EY032532) to S.G.W., SUNY Startup Funds to S.G.W, SUNY Graduate Assistantship to D.L., S.K., and S.C., NEI T35 to A.M.R. Generative AI (ChatGPT, OpenAI) was used to assist with the generation of the summary illustration. Grammarly and ChatGPT were used for language editing and refinement of the manuscript text. All AI content was critically reviewed and approved by the authors.

