## Supplementary material for "Disabling Müller Glia Preserves Retinal Function After Retinal Injury": Larbi 2026_Supplement revision

Supplement Material

Larbi et al., 2026

**Table S1: Genotyping Primers**

| Gene name | Forward sequence (5' to 3') | Reverse sequence (3' to 5') |
|---|---|---|
| **Rlbp1Cre transgene** | CAA GTG TGA GAG ACA GCA TTG | TCC TTA GCG CCG TAA ATC AA |
| **Glast1Cre transgene** | ACA ATC TGG CCT GCT ACCAAA GC | CCA GTG AAA CAG CAT TGCTGT C |
| **Ascl1Cre wildtype** | TCC AAC GAC TTG AAC TCT ATG G | CCA GGA CTC AAT ACG CAG GG |
| **Ascl1Cre mutant** | AAC TTT CCT CCG GGG CTC GTT TC | CGC CTG GCG ATC CCT GAA CAT G |
| **tdTomato wildtype** | AAG GGA GCT GCA GTG GAG TA | CCG AAA ATC TGT GGG AAG TC |
| **tdTomato mutant** | CTG TTC CTG TAC GGC ATG G | GGC ATT AAA GCA GCG TAT CC |
| **Dicer** | CCTGACAGTGACGGTCCAAAG | CATGACTCTTCAACTCAAACT |
| **RPE65** | CAC TGT GGT CTC TGC TAT CTT C | GGT GCA GTT CCA CTT CAG TT |

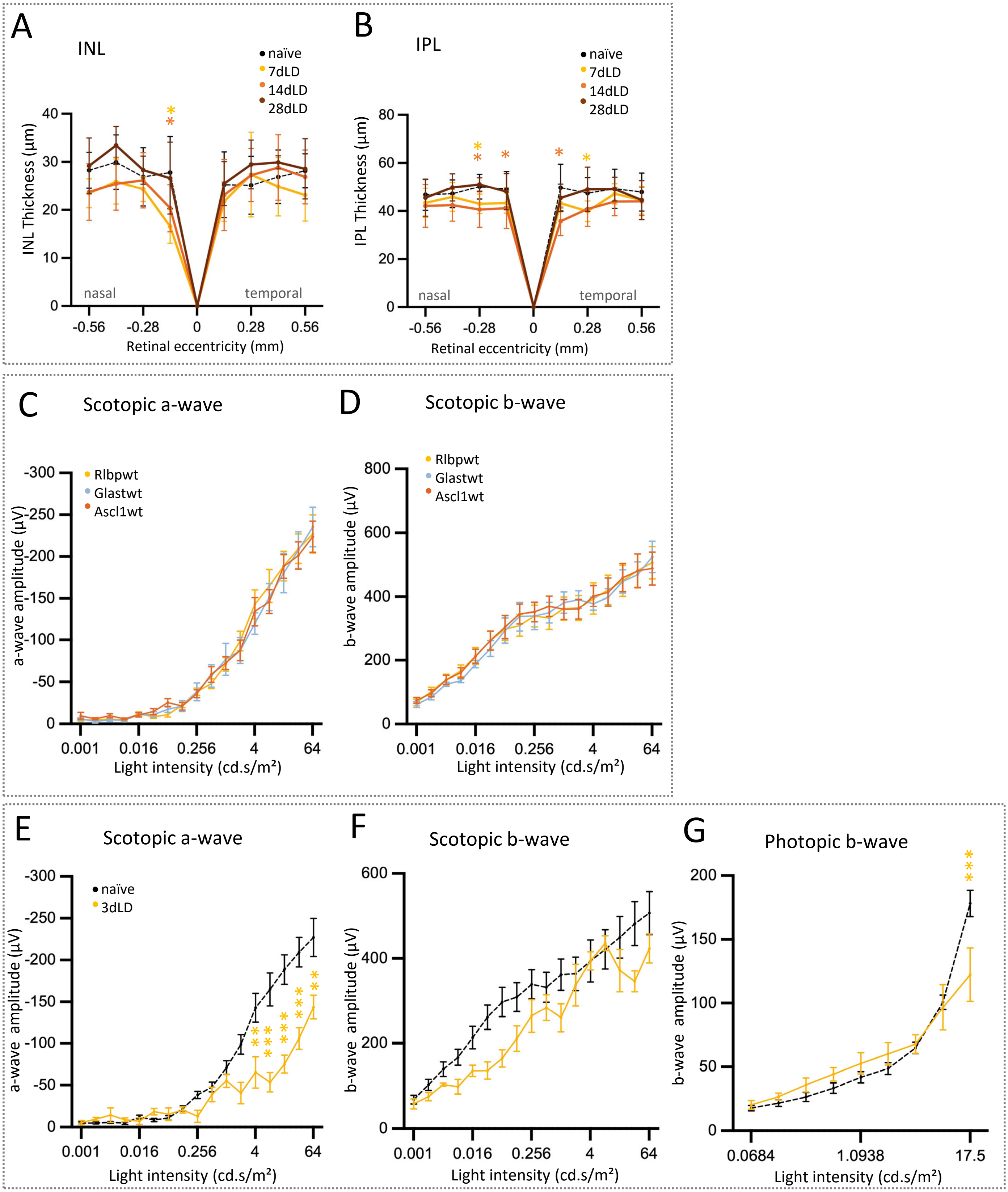


**Figure S1: OCT and ERG of undamaged and damaged retinas. A-B:** Spider plots (nasal-temporal axis) and time course (averaged nasal-temporal and superior-inferior axis) of the inner nuclear layer (A) and inner plexiform layer (B) of undamaged (n=11) or light-damaged mice, 7d (n=8), 14d (n=12), or 28dpLD (n=6). Two-way mixed-effects model with Dunnett's multiple comparisons correction comparing all timepoints to the naïve control, *: p≤0.05. **C-G:** Full-field ERG recordings showing scotopic a-wave amplitudes (C, E), scotopic b-wave amplitudes (D, F), and photopic b-wave amplitudes (G), in undamaged 3-month Rlbp-Cre (n=12), 3-month Glast-cre (n=5), 2-month Ascl1-Cre-driven reporter wildtype mice (n=10) as, as well as 3 days after light damage (3dLD, n=4, E-G). OCT: Mean ± S.D.; ERG: Mean ± S.E.M. Two-way ANOVA with Tukey's multiple comparisons correction (C,D) and two-way ANOVA with Bonferroni's multiple comparisons correction, **: p≤0.01, ***: p≤0.001 (E-G).

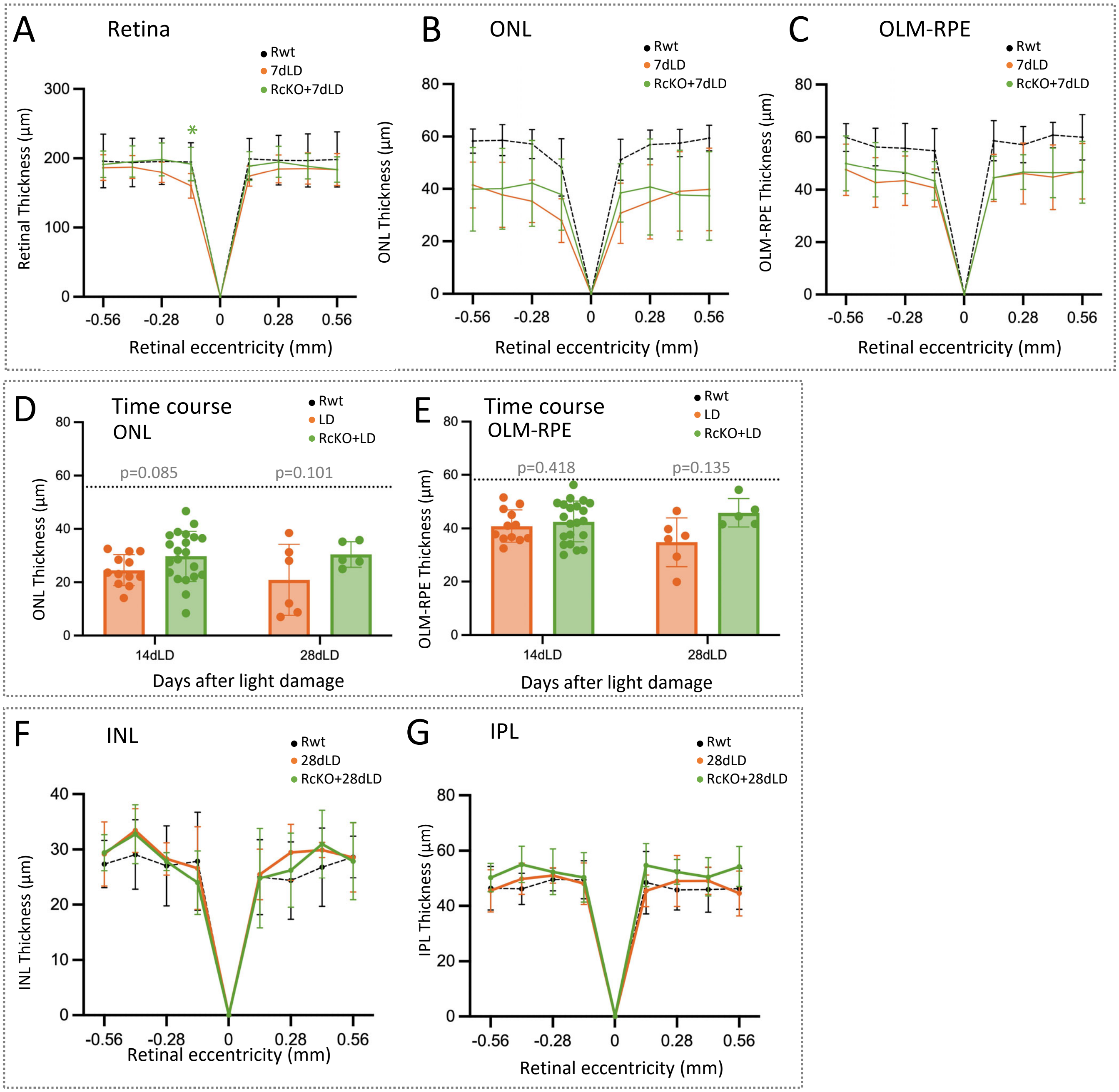


**Figure S2: OCT of light damaged Rlbp-Cre:Dicer-cKO mice. A-C:** Spider plots (nasal-temporal axis) of the total retinal thickness (A), outer nuclear layer thickness (B), and OLM-RPE thickness (C) of undamaged Rlbp-Cre wildtype mice (Rwt, n=7), light-damaged Rlbp-Cre:Dicer-cKO mice (RcKO+LD) 7dpLD or light-damaged wildtypes (LD) 7dpLD (n=8). Two-way ANOVA with Tukey's multiple comparisons correction for comparisons 7dLD vs. RcKO+7dLD, *: p≤0.05. **D-E:** Time course of averaged thickness of the nasal-temporal and superior-inferior axis of the outer nuclear layer (ONL, D) and the outer limiting membrane/retinal pigment epithelium (OLM-RPE, E) of light-damaged Rlbp-Cre:Dicer-cKO mice (RcKO+LD) 14d (n=21), or 28d pLD (n=5) or light damaged wildtypes (LD),14d (n=12), or 28d pLD (n=6). Undamaged wildtype baseline is shown as dotted line. Linear mixed-effects model with post hoc comparisons using estimated marginal means and Bonferroni correction for multiple comparisons, p-values are given in the graphs. **F-G:** Spider plots (nasal-temporal axis) of the inner nuclear layer (F) and inner plexiform layer (G) of Rlbp-Cre:Dicer-cKO mice (RcKO+LD) 28dpLD (n=5) or light-damaged wildtypes (LD) 28dpLD (n=6). Two-way ANOVA with Tukey's multiple comparisons correction for comparisons 28dLD vs RcKO+28dLD, *: p≤0.05, **: p≤0.01, ***: p≤0.001.

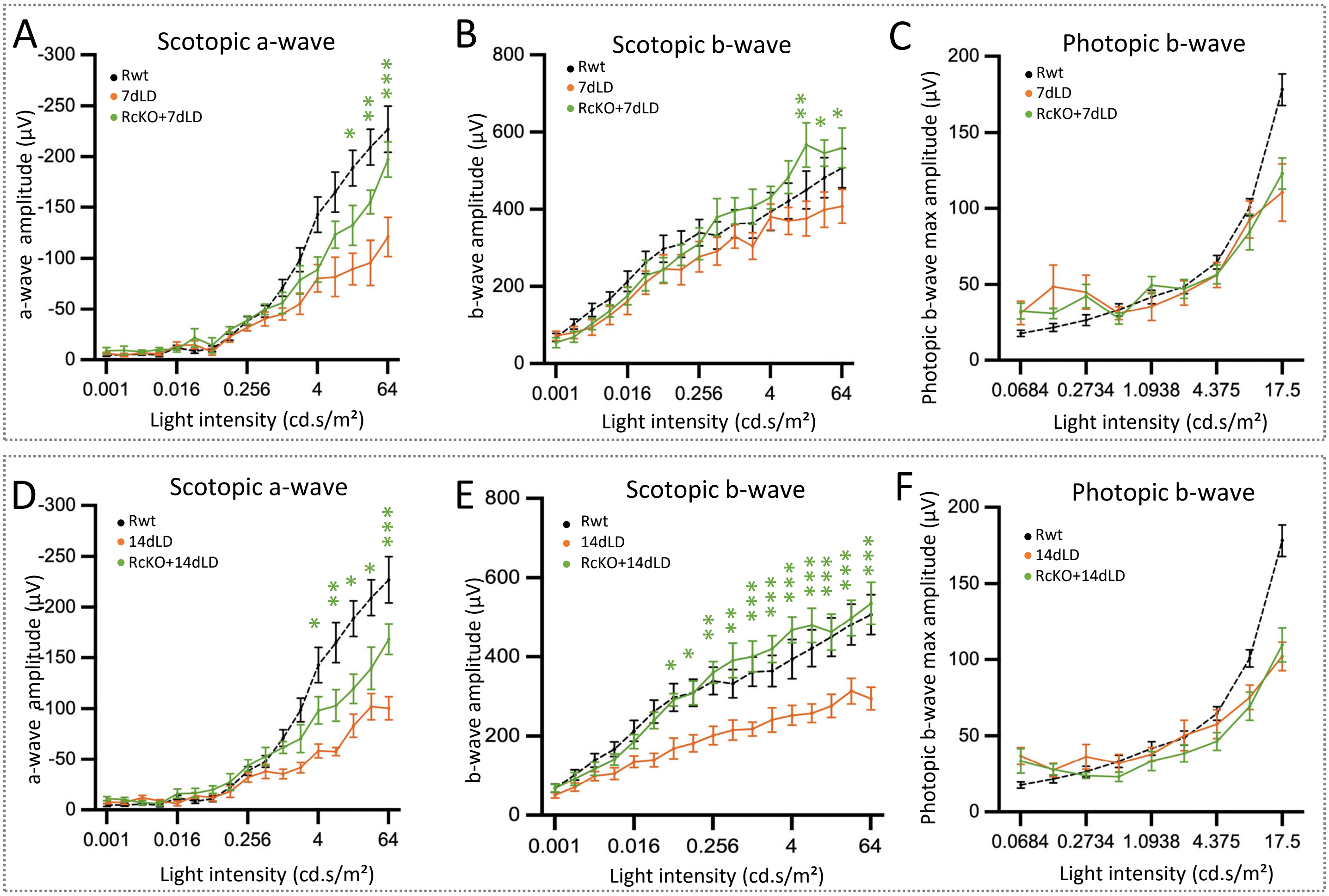


**Figure S3: ERG of light-damaged Rlbp-Cre:Dicer-cKO mice. A-F:** Full-field ERG recordings showing scotopic a-wave (A, D), scotopic b-wave (B, E), as well as photopic b-wave amplitudes (C, F) of undamaged Rlbp-Cre wildtypes (Rwt, n=12), light-damaged Rlbp-Cre:Dicer-cKO mice (RcKO+LD), 7d (n=7), 14d (n=8) and light-damaged mice (LD), 7d (n=7), 14d (n=7). Mean ± S.E.M. Two-way ANOVA with Tukey's multiple comparisons correction; comparisons displayed are 7dLD vs RcKO+7dLD (A,B,C), 14dLD vs RcKO+14dLD (D-F), *: p≤0.05, **: p≤0.01, ***: p≤0.001.

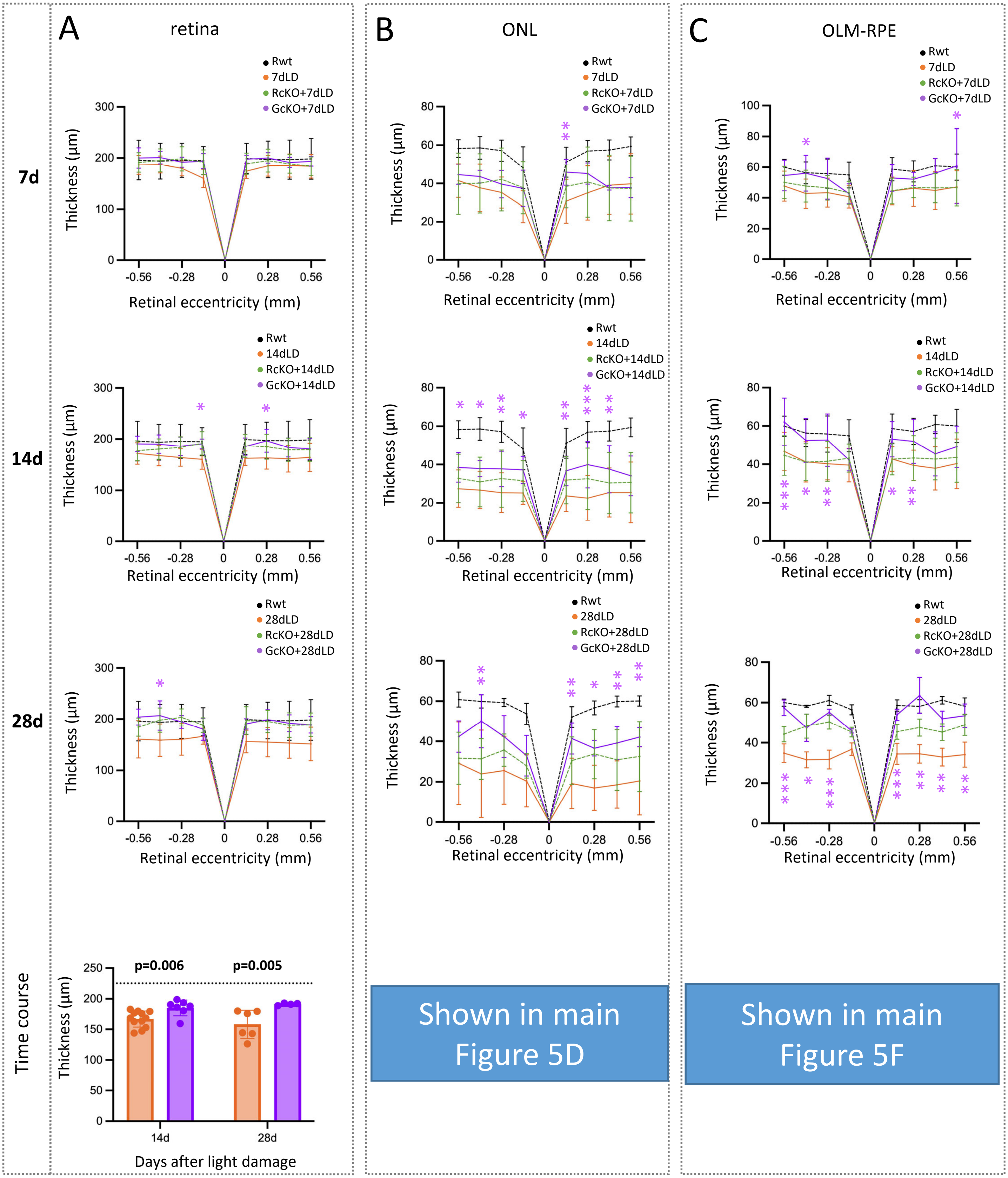


**Figure S4: OCT of light-damaged Glast-Cre: Dicer-cKO mice. A-C:** Spider plots (nasal-temporal axis) and time courses (averaged nasal-temporal and superior-inferior axis) of the thickness (diameter, µm) of the total retina (A), the outer nuclear layer (ONL, B), and the outer limiting membrane/retinal pigment epithelium (OLM-RPE, C) of undamaged mixed wildtypes (n=12), light-damaged Glast-Cre: Dicer-cKO (GcKO+LD) 7d (n=5), 14d (n=7), or 28dpLD (n=4), light-damaged Rlbp-Cre: Dicer-cKO (RcKO+LD) 7d (n=12), 14d (n=21), or 28dpLD (n=5), and for light-damaged wildtypes (LD) 7d (n=8), 14d (n=12), or 28dpLD (n=6). Mean ± S.D. Single timepoints were analyzed by two-way ANOVA with Tukey's multiple comparisons correction; comparisons displayed are LD vs. GcKO+LD at the indicated timepoint, p≤0.05, **: p≤0.01, ***: p≤0.001. Time course data were analyzed using linear mixed-effects model with post hoc comparisons using estimated marginal means and Bonferroni correction for multiple comparisons, p-values are given in the graphs. Undamaged wildtype values serve only as a reference/baseline; Rlbp-Cre: Dicer-cKO data are plotted only for reference (dotted lines).

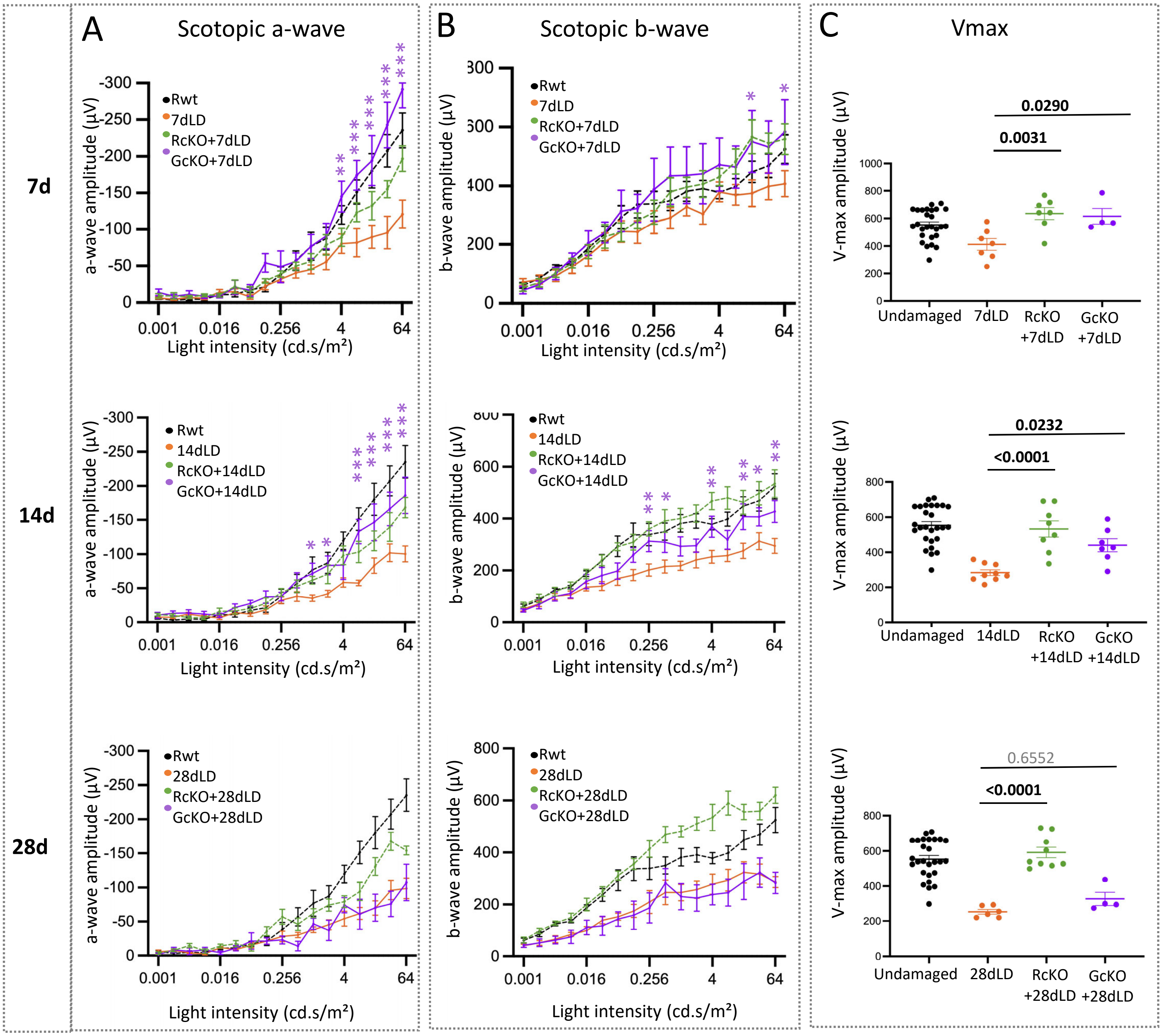


**Figure S5: ERG of light-damaged Glast-Cre:Dicer-cKO mice. A-C:** Full-field ERG recordings showing scotopic a-wave amplitudes and corresponding time line of the maximum averaged amplitudes (A), scotopic b-wave amplitudes (B) and estimated saturated amplitudes (Vmax, responsiveness) using the Naka-Rushton equation (C) of undamaged mixed wildtypes (n=17), light-damaged Glast-Cre:Dicer-cKO (GcKO+LD) 7d (n=4), 14d (n=7), or 28dpLD (n=4), light-damaged Rlbp-Cre:Dicer-cKO (RcKO+LD) 7d (n=7), 14d (n=8), or 28dpLD (n=5), and for light-damaged wildtypes 7d (n=7), 14d (n=9), or 28dpLD (n=6). Mean ± S.E.M. Single timepoints were analyzed by two-way ANOVA with Tukey's multiple comparisons correction; comparisons displayed are LD vs. GcKO+LD at the indicated timepoint, $p \leq 0.05$, **: $p \leq 0.01$, ***: $p \leq 0.001$. Time course data were analyzed using linear mixed-effects model with post hoc comparisons using estimated marginal means and Bonferroni correction for multiple comparisons, p-values are given in the graphs. Undamaged wildtype values serve only as a reference/baseline; Rlbp-Cre: Dicer-cKO data are plotted only for reference (dotted lines).

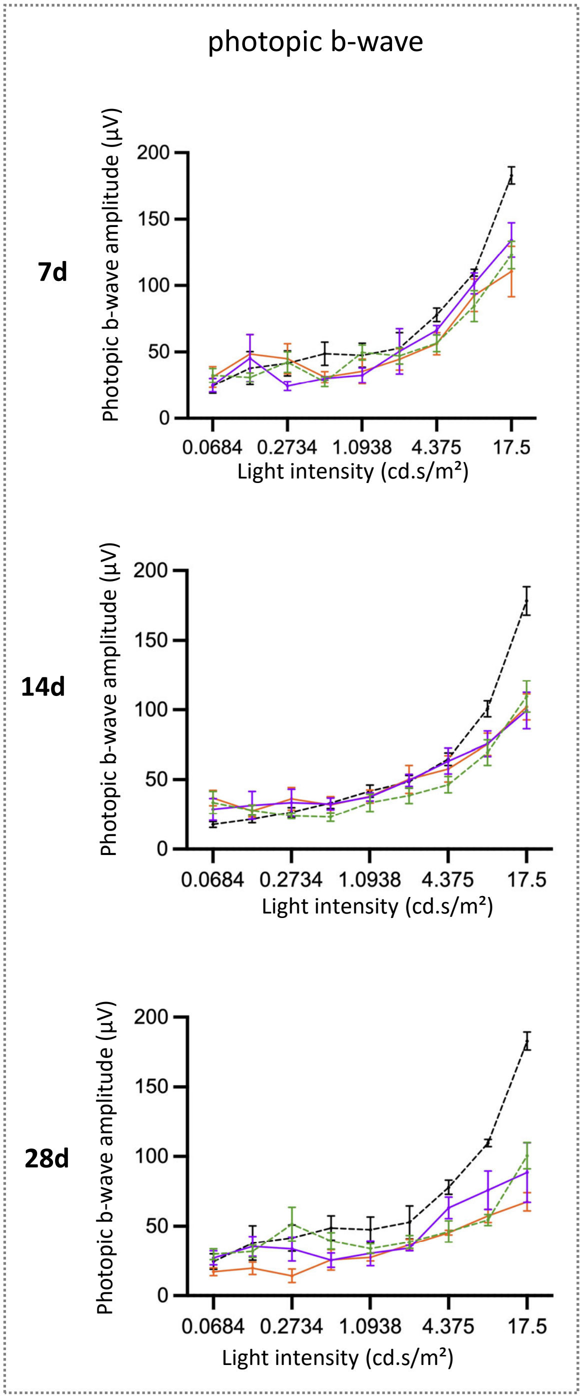


**Figure S6: Photopic b-wave amplitudes of light-damaged Glast-Cre:Dicer-cKO mice.** b-wave amplitudes per time point and corresponding time line of undamaged mixed wildtypes (n=15), light-damaged Glast-Cre:Dicer-cKO (GcKO+LD) 7d (n=4), 14d (n=7), or 28dpLD (n=4), light-damaged Rlbp-Cre:Dicer-cKO (RcKO+LD) 7d (n=7), 14d (n=8), or 28dpLD (n=5), and for light-damaged wildtypes 7d (n=7), 14d (n=9), or 28dpLD (n=6). Mean ± S.E.M.. Single timepoints were analyzed by two-way ANOVA with Tukey's multiple comparisons correction; comparisons displayed are LD vs. GcKO+LD at the indicated timepoint, p≤0.05, **: p≤0.01, ***: p≤0.001. Time course data were analyzed using linear mixed-effects model with post hoc comparisons using estimated marginal means and Bonferroni correction for multiple comparisons, p-values are given in the graphs. Undamaged wildtype values serve only as a reference/baseline; Rlbp-Cre: Dicer-cKO data are plotted only for reference (dotted lines).

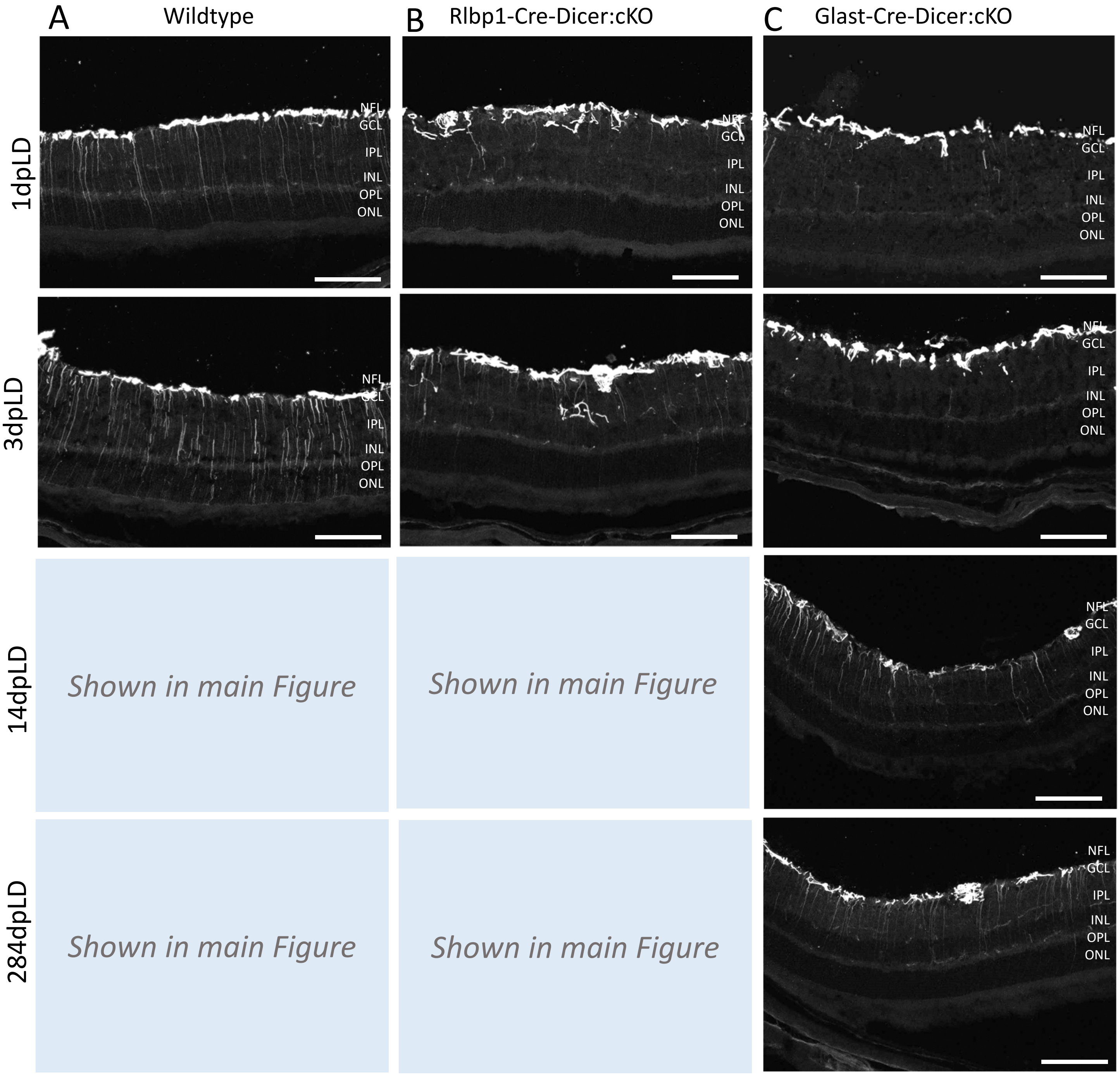


**Figure S7: GFAP immunoreactivity at early light-damage early stages. A-I:** Immunofluorescent labeling with antibodies against GFAP of retinal cross sections of representative damage areas of wildtype (panel A), Rlbp1-Cre: Dicer-cKO (panel B) or Glast-Cre: Dicer-cKO mice (panel C), 1, 3, 14, or 28 days after light damage (dpLD). Scale bars 100 µm. NFL: Nerve Fiber Layer, GCL: ganglion cell layer, IPL: inner plexiform layer, INL: inner nuclear layer, OPL: outer plexiform layer, ONL: outer nuclear layer, OLM: outer limiting membrane.
